# Ontogenetic changes in sensory gene expression in *Bicyclus anynana* butterflies

**DOI:** 10.1101/2020.09.15.298133

**Authors:** David A. Ernst, Erica L. Westerman

## Abstract

Animal behavior is largely driven by the information that animals are able to extract and process from their environment. However, the function and organization of sensory systems often change throughout ontogeny, particularly in animals that undergo indirect development. As an initial step toward investigating these ontogenetic changes at the molecular level, we characterized the sensory gene repertoire and examined the expression profiles of genes linked to vision and chemosensation in two life stages of an insect that undertakes a dramatic metamorphosis, the butterfly *Bicyclus anynana*. Using RNA-seq, we compared gene expression in the heads of late fifth instar larvae and newly-eclosed adults that were reared under identical conditions. Over 50% of all expressed genes were differentially expressed between the two developmental stages, with 4,046 genes upregulated in larval heads and 4,402 genes upregulated in adult heads. In larvae, upregulated vision-related genes were biased toward those involved with eye development, while phototransduction genes dominated the vision genes that were upregulated in adults. Moreover, the majority of the chemosensory genes we identified in the *B. anynana* genome were differentially expressed between larvae and adults, several of which share homology with genes linked to pheromone detection, host plant recognition, and foraging in other species of Lepidoptera. These results reveal promising candidates for furthering our understanding of sensory processing and behavior in the disparate developmental stages of butterflies and other animals that undergo metamorphosis.

## Introduction

The environment is teeming with information, and the ability to perceive and process this information is critical in shaping the behavior of all animals. Of the various sensory modalities, vision and chemoreception play integral roles in survival and reproduction, including the detection of food sources (Nevitt, 2008), predator avoidance (Fischer, Oberhummer, Cunha-Saraiva, Gerber, & Taborsky, 2017), and locating potential mates (Shine, Webb, Lane, & Mason, 2005). Moreover, both senses are known to drive assortative mating and speciation processes. Visual cues, such as ornaments (Robertson & Monteiro, 2005), coloration (Bakker & Mundwiler, 1994), and mating displays (Wiley, 1973), influence mate choice behaviors and sexual selection in a diverse range of species. Similarly, chemical signals, such as pheromones and cuticular hydrocarbons, have been found to be involved with prezygotic isolation in a wide variety of taxa, ranging from insects (Coyne, Crittenden, & Mah, 1994; Pélozuelo, Meusnier, Audiot, Bourguet, & Ponsard, 2007; Raymond, Searle, & Douglas, 2001) and annelids (Sutton et al., 2005) to mammals (Laukaitis, Critser, & Karn, 1997; Moore, 1965; Nevo, Bodmer, & Heth, 1976).

Despite the significant roles that visual and chemical cues play in animal behavior and sexual selection, considerable morphological differences often exist for sensory structures throughout ontogeny. This is particularly apparent in animals that undergo metamorphosis from larva to adult life stages, such as holometabolous insects (Truman & Riddiford, 1999), crustaceans (Passano, 1961), and many fishes (McMenamin & Parichy, 2013). For instance, in butterflies, the visual organs of the larval stage typically consist of up to six stemmata per eye, each with a lens and seven photoreceptors that form a tiered rhabdom (Gilbert, 1994), compared to the much more complex adult compound eyes, which consist of hundreds of tightly-packed ommatidia, each containing a facet lens and rhabdom composed of nine photoreceptors (Arikawa, 2017). These differences are likely in part due to the different ecological niches that each stage fills; larvae typically reside and forage on host plants until pupation, while adults adopt an aerial lifestyle and are mainly focused on finding a mate and reproducing (Truman & Riddiford, 1999).

While differences in sensory organ morphology and the behavior of animals that undergo metamorphosis are often readily apparent throughout ontogeny, we still have much to learn about the functional and organizational differences in the sensory systems of pre- and post-metamorphosis life stages, especially at the molecular level. Perhaps one of the most promising taxa in which to dissect these differences is the exceptionally diverse Insecta, which is estimated to consist of 5.5 million species (Stork, 2018). Indeed, much of what we know about the molecular mechanisms underlying vision and chemosensation has been derived from work on the common fruit fly, *Drosophila melanogaster* (see Hardie, 2001 and Benton, 2008 for review).

Phototransduction in insects is accomplished in the eye through absorption of light by a visual pigment (rhodopsin), which triggers an enzymatic cascade that ultimately leads to depolarization of photoreceptor cells (Hardie, 2001). The perception of different wavelengths of light is dependent upon opsin structure, with peak sensitivities spanning the visible light spectrum and beyond (Terakita, 2005). By contrast, chemosensation in insects occurs at the olfactory sensilla (the sensory structures involved with smell) typically found on head structures, such as the maxillary palps and antennae, and the gustatory sensilla (the sensory structures involved with taste), which are found throughout the insect body, including on the mouthparts, wings, and legs (Briscoe et al., 2013; Anupama Dahanukar, Hallem, & Carlson, 2005). Odorants are bound by odorant binding proteins (OBPs) or chemosensory proteins (CSPs) and transported through the sensillar lymph to membrane-bound receptors located on the dendrites of olfactory sensory neurons (OSNs) or gustatory sensory neurons (GSNs) (Pelosi, Calvello, & Ban, 2005; Vogt & Riddiford, 1981).

There are three types of chemoreceptors on chemosensory neurons that are involved with the detection of chemical stimuli from the external environment in insects: odorant receptors (ORs), gustatory receptors (GRs), and ionotropic receptors (IRs). ORs are the foundation of olfaction and are known to selectively detect a diversity of volatile compounds (Ha & Smith, 2009; Hansson & Stensmyr, 2011; Sato et al., 2008). In combination with a co-receptor (*Orco*) and sensory neuron membrane proteins (SNMPs), some ORs have also been found to be involved with the detection of sex pheromones (Benton, Vannice, & Vosshall, 2007; Zhang et al., 2020). Insect GRs belong to the same superfamily as ORs (Clyne, Warr, & Carlson, 2000) but are primarily involved with tasting bitter compounds (Lee, Moon, & Montell, 2009; Moon, Köttgen, Jiao, Xu, & Montell, 2006; Weiss, Dahanukar, Kwon, Banerjee, & Carlson, 2011), sugars (Chyb, Dahanukar, Wickens, & Carlson, 2003; Dahanukar, Foster, Van der Goes van Naters, & Carlson, 2001; Slone, Daniels, & Amrein, 2007), and CO2 (Jones, Cayirlioglu, Grunwald Kadow, & Vosshall, 2007; Kwon, Dahanukar, Weiss, & Carlson, 2007). Finally, IRs, which are primitive chemoreceptor proteins that evolved from ionotropic glutamate receptors (iGluRs), are known to be involved with both olfaction and gustation, primarily sensing amines, acids, salt, and pheromones (Briscoe et al., 2013; Koh et al., 2014; Rytz, Croset, & Benton, 2013; Zhang, Ni, & Montell, 2013).

Recent work in adult Lepidoptera has focused on elucidating the underpinnings of phototransduction (Macias-Muñoz, Rangel Olguin, Briscoe, & Li, 2019) and chemoreception (Briscoe et al., 2013; W. Liu, Jiang, Cao, Yang, & Wang, 2018; Vogt, Große-Wilde, & Zhou, 2015; Zhang et al., 2020), providing a foundation for investigating how sensory systems vary throughout development in an insect order known for its radical metamorphosis. The squinting bush brown butterfly, *Bicyclus anynana*, is an ideal model to address this question, as it has rapidly become a fruitful model system for studying development, evolution, and phenotypic plasticity (Brakefield & Reitsma, 1991; Koch, Brakefield, & Kesbeke, 1996; Kooi & Brakefield, 1999; Prudic, Jeon, Cao, & Monteiro, 2011). Of particular interest, these butterflies rely heavily on visual and chemical cues for mate choice; mates are selected based on the quality of ultraviolet-reflective eyespot pupils and male-specific pheromones (Costanzo & Monteiro, 2007; Robertson & Monteiro, 2005; Westerman & Monteiro, 2013). In addition, previous work has identified differences in the visual systems of male and female adults of different seasonal phenotypes, including differences in eye size, facet lens area, facet number per eye, and opsin and eye development gene expression (Everett, Tong, Briscoe, & Monteiro, 2012; Macias-Munoz, Smith, Monteiro, & Briscoe, 2016). Importantly, numerous molecular resources are available for this species, including a reference genome assembly (Nowell et al., 2017), making this *B. anynana* amenable to genetic and genomic studies.

Here, we characterized the sensory gene repertoire in *B. anynana*, including genes known to be linked to vision and chemosensation. Specifically, we first identified vision genes in the *B. anynana* genome involved with phototransduction, eye pigmentation, and eye development, as well as six distinct families of chemosensory genes, consisting of OBPs, CSPs, ORs, IRs, GRs, and SNMPs. We then compared the expression patterns of these sensory genes in the heads of two developmental stages: late fifth instar larvae and newly-eclosed adult butterflies. Additionally, we investigated the expression of genes that are known to be involved with wing patterning in *B. anynana* and other butterflies, which are hypothesized to potentially drive speciation and assortative mating by linking wing pattern traits to preference for those traits (Kronforst et al., 2006; Merrill et al., 2019; Westerman, 2019).

We predicted that genes directly involved with visual processes (e.g., phototransduction) would be upregulated in the adult phenotype, given the much greater level of complexity of adult compound eyes compared to the relative simplicity of larval stemmata. In addition, we hypothesized that larvae might not express the ultraviolet-sensitive opsin that is critical to eyespot evaluation during mate choice in adults. For chemosensory genes, we predicted that genes involved with pheromone and fruit detection would be upregulated in adults, as this stage participates in numerous reproductive behaviors and must locate a food source (i.e., ripe or rotting fruit) separate from the host plant. By contrast, we hypothesized that chemosensory genes linked to host plant recognition and foraging behavior would be upregulated in larvae, given that feeding is the dominant behavior during this stage of development. Finally, we aimed to elucidate candidate visual and chemosensory genes in the adult and larval phenotypes for future investigation into the sensory ecology of these disparate life stages.

## Materials and Methods

### Animals

*Bicyclus anynana*, a Nymphalid butterfly native to subtropical Africa, has been maintained in laboratory colonies since 1988. All animals used in this study are descendants of an original population established in Leiden, Netherlands from 80 gravid females that were collected in Malawi (Brakefield & Reitsma, 1991). The population at the University of Arkansas was established via the transfer of ~1,000 eggs from a population in Singapore to Fayetteville, AR, USA in spring, 2017. All animals were reared in a climate-controlled, USDA-APHIS approved (Permit # P526P-17-00343) greenhouse facility, which was maintained at approximately 27°C, 70% relative humidity, and under a 13:11h light:dark photoperiod to induce the wet season phenotype in this species (Brakefield & Larsen, 1984).

### Experimental design and tissue collection

All experiments took place between January and April 2019. Four separate families were created by pairing one three-day-old naïve male and one three-day-old naïve female together in a small mesh cage (31.8 cm × 31.8 cm × 31.8 cm) at 8:00 am for at least three hours to ensure that copulation occurred. After visual confirmation that the pair had copulated, the female was removed from the mating cage and isolated in a new large mesh cage (39.9 cm × 39.9 cm × 59.9 cm) containing a corn plant on which to lay eggs and slice of moistened banana for food. Each female was then given seven days to lay fertilized eggs on the provided corn plant, after which the egg-laden corn plant was transferred to a new small mesh cage (31.8 cm × 31.8 cm × 31.8 cm).

Upon hatching, larvae were reared in their family-specific cages under identical conditions and were fed corn plants *ad libitum*. To ensure that all four families experienced the same environmental conditions within the greenhouse and to control for any potential unforeseen confounding variables associated with cage location, the physical position of each cage was alternated daily. Upon the morning of reaching the late fifth instar stage, which was determined by the stark change in color from tan/brown to green (Fig. 1A), a subset of the larvae from each family was sacrificed by decapitation with RNase-free scissors. A second subset from each family was allowed to pupate, and newly-eclosed adults (Fig. 1B) were sacrificed by decapitation on the morning of emergence. All heads were immediately transferred into RNase-free, low binding 1.5 ml microcentrifuge tubes (Biotix, San Diego, CA, USA), flash-frozen in liquid nitrogen, and transported to the lab for storage at −80°C until they were processed (Table S1).

**Fig. 1:**
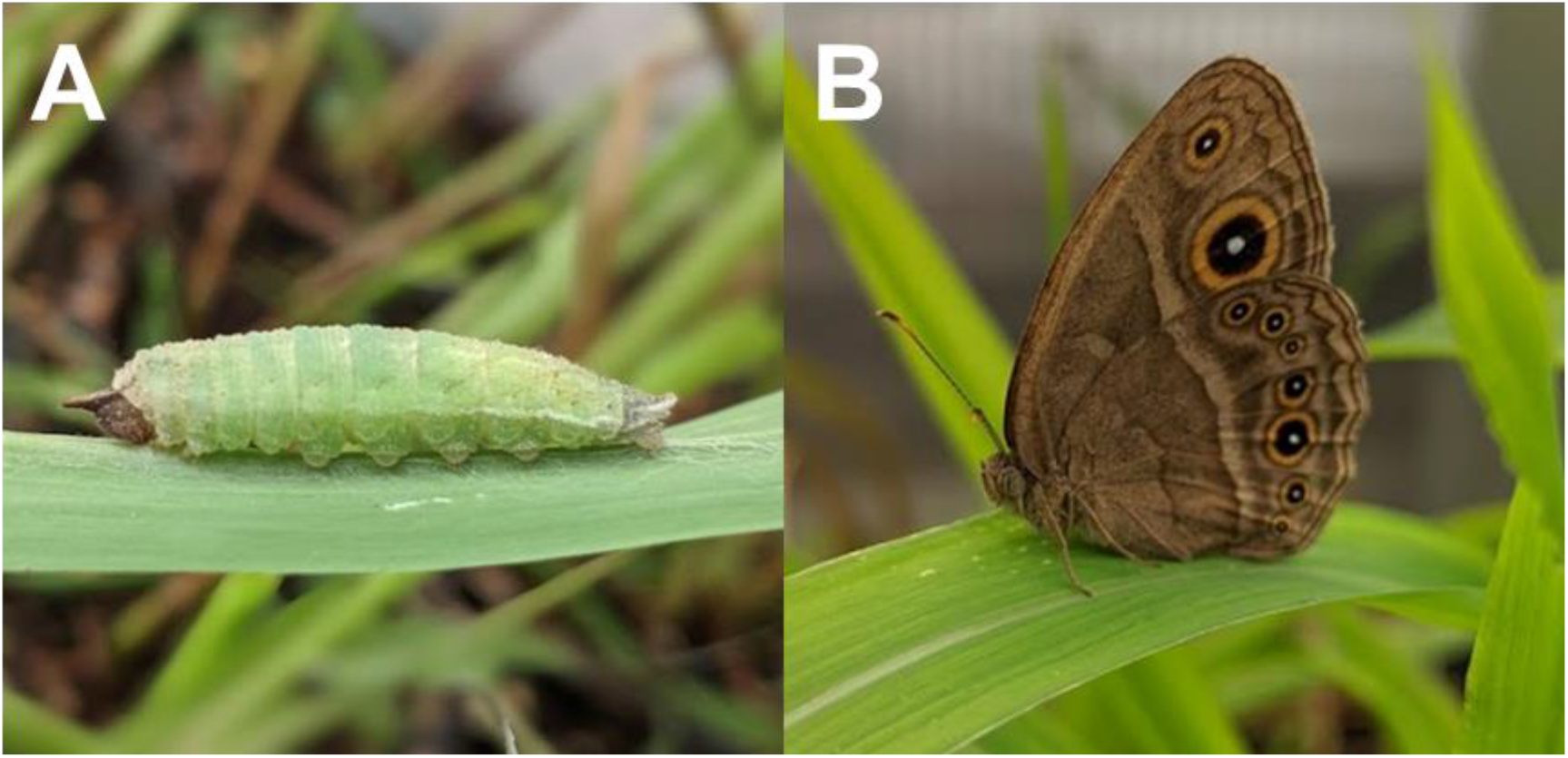
Developmental stages of *B. anynana*. (A) Late fifth instar larva. (B) Newly-eclosed adult *B. anynana* (not to scale).

### RNA extraction, library preparation, and sequencing

Each frozen head was immersed in pre-chilled RNAlater-ICE (Ambion; Austin, TX, USA) and incubated at −20°C for approximately 16 h prior to tissue processing. After this incubation period, heads were transferred to a dissecting dish filled with RNAlater-ICE, and all residual thoracic tissue was carefully removed with forceps under a dissecting microscope (Zeiss Stemi 508; Jena, Germany), leaving only head tissue. Isolated heads (which included antennae and mouthparts) were then disrupted in lysis buffer with an RNase-free, disposable pestle, and small (<200 nt) and large RNA (>200 nt) were extracted in separate fractions using the NucleoSpin^®^ miRNA kit (Macherey-Nagel; Düren, Germany) following the manufacturer’s recommended protocols. RNA purity, concentration, and integrity for each sample were subsequently determined using a NanoDrop 2000 (Thermo Fisher Scientific; Waltham, MA, USA) and TapeStation 2200 (Agilent; Santa Clara, CA, USA).

After confirmation of RNA quality and quantity, cDNA libraries were prepared using 500 ng of large RNA as input for the KAPA mRNA HyperPrep Kit (Kapa Biosystems; Wilmington, MA, USA) combined with the KAPA Unique Dual-Indexed Adapter Kit (Kapa Biosystems; Wilmington, MA, USA). The quality of each cDNA library was subsequently verified using a TapeStation 2200 (Agilent; Santa Clara, CA, USA). All libraries were then shipped on dry ice to the University of Chicago Genomics Facility for secondary quality assessment on a 5300 Fragment Analyzer (Agilent; Santa Clara, CA, USA), and 50 base pair (bp) single-end (SE) sequencing was performed on a single lane of a HiSeq 4000 (Illumina; San Diego, CA, USA).

### Functional annotation

Blast2GO v5.2.5 (Conesa et al., 2005) was used to conduct a *de novo* functional annotation of all genes in the most current *B. anynana* reference genome (v1.2; Nowell et al., 2017). First, we used BLASTx v2.6.0+ (Altschul, Gish, Miller, Myers, & Lipman, 1990) to search the NCBI ‘nr’ protein database (www.ncbi.nlm.nih.gov/refseq) and collected the top 10 hits with an e-value <10^-3^. These results were then uploaded into Blast2GO, and further functional classification was performed using the InterProScan (Jones et al., 2014) function within Blast2GO. Finally, the “Mapping” and “Annotation” steps in Blast2GO were performed using the default parameters, and the resulting functional annotation table was exported.

### Differential gene expression analysis

Prior to expression quantification, the quality of the raw reads was assessed using FastQC v0.11.8 (https://www.bioinformatics.babraham.ac.uk/projects/fastqc/), and Illumina adapter sequences were trimmed using Trimmomatic v0.38 (Bolger, Lohse, & Usadel, 2014). Trimmed reads were then aligned to the *B. anynana* reference genome (v1.2) using STAR v2.7.1a (Dobin et al., 2013). Reads were then quantified using the ‘htseq-count’ command in the HTSeq v0.11.2 Python package (Anders, Pyl, & Huber, 2015). Differential gene expression analysis was conducted using the DESeq2 v1.24.0 package (Love, Huber, & Anders, 2014) in R (Version 3.6.2, R Foundation for Statistical Computing, Vienna, Austria). The generalized linear model:

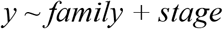

was fit to each gene using a negative binomial distribution, where *y* denotes the response variable (expression), *family* denotes the family to which each individual belongs (family 1-4), and *stage* denotes the life stage of each individual (larva or adult). Using this design enabled us to contrast the effect of *stage* while controlling for differences in expression associated with lineage. Genes with a total read alignment count <10 were filtered and not included in the differential expression analysis. Gene expression was calculated as the binary log of the expression fold change (log_2_FC), and the *apeglm* method was used for log_2_FC shrinkage (Zhu, Ibrahim, & Love, 2019). Finally, genes with a false discovery rate (FDR; Benjamini & Hochberg, 1995) <0.05 were retained for downstream analysis.

### Gene ontology enrichment analyses

For further characterization, the set of differentially expressed genes was split into genes that showed increased expression in adults (log_2_FC >0) and those that showed increased expression in larvae (log_2_FC <0). The Fisher’s Exact Test function in Blast2GO was used to test for GO term enrichment in each of the differentially expressed gene sets separately with all genes in the expression set as the reference set, and only GO terms with an FDR <0.05 were considered significantly enriched. The list of enriched GO terms was then reduced to the most specific terms for visualization. Additionally, the reduced lists of enriched GO terms were processed using REVIGO (http://revigo.irb.hr/; Supek, Bošnjak, Škunca, & Šmuc, 2011), which further eliminated redundancy and organized GO terms into treemaps consisting of related superclusters.

### Identification of visual genes

To identify differentially expressed genes involved with vision (i.e., phototransduction, eye pigment, and eye development) in *B. anynana*, we first collected the coding sequences (CDS) of 74 putative *Heliconius melpomene* phototransduction genes from Macias-Muñoz et al. (2019) and the protein sequences of 200 *D. melanogaster* phototransduction, eye pigment, and eye development genes compiled by Macias-Munoz et al. (2016). We then used BLASTX and BLASTP (BLAST v2.2.30+; Altschul et al., 1990) to query these sequences against the *B. anynana* reference genome proteins and identify homologs. Homologs were determined based on hits with an E-value <1E-10. Finally, we manually searched the Blast2GO annotation descriptions, best blast hits, and GO annotations for terms linked to vision, including: “eye,” “ommatidia,” “ommatidium,” “opsin,” “photoreceptor,” “phototransduction,” “retina,” and “visual.”

### Identification of chemosensory genes

To identify differentially expressed genes involved with chemosensation in *B. anynana*, we collected 273 lepidopteran and *D. melanogaster* odorant binding protein (OBP) protein sequences from Vogt et al. (2015), 34 *H. melpomene* chemosensory protein (CSP) protein sequences from Dasmahapatra et al. (2012), 70 *H. melpomene* olfactory receptor (OR) protein sequences from Dasmahapatra et al. (2012), 31 *B. anynana* IR sequences from Liu et al. (2018), 73 *H. melpomene* gustatory receptor (GR) protein sequences from Briscoe et al. (2013), and 33 lepidopteran sensory neuron membrane protein (SNMP) sequences from Zhang et al. (2020). These sequences were then queried against the *B. anynana* reference genome with BLASTX or BLASTP to identify putative homologs. With the exception of the previously identified IR sequences in *B. anynana*, all hits with an E-value <1E-10 were further screened for conserved protein domains specific to each gene family using CD-Search (Marchler-Bauer & Bryant, 2004). Specifically, sequences with hits for the following domains were retained: OBPs, either pfam01395 (PBP/GOBP family) *or* smart00708 (Insect pheromone/odorant binding protein domains); CSPs, pfam03392 (Insect pheromone-binding family, A10/OS-D); ORs, either pfam02949 (7tm Odorant receptor) *or* pfam08395 (7tm Chemosensory receptor); GRs, pfam08395 (7tm Chemosensory receptor); and SNMPs, pfam01130 (CD36 family). Because the CD36 superfamily common to SNMPs consists of three different protein families, only one of which includes SNMPs (Vogt et al., 2009), we filtered the final putative SNMP sequences by only retaining those that were also annotated as SNMPs in our functional annotation. Finally, we performed a manual search of the Blast2GO functional annotation to identify any additional putative OBP, CSP, OR, IR, GR, and SNMP genes. Specifically, we searched the Blast2GO descriptions, best blast hits, and GO annotations for key terms, including: “chemoreceptor,” “chemosensory,” “gustatory,” “odor,” “olfaction,” “olfactory,” “pheromone,” and “smell” and subjected any putative chemosensory genes to the conserved protein domain filtration described above.

### Identification of wing patterning genes

To further explore genes that might be relevant to sensory processing during mate choice, we also manually searched our *de novo* functional annotation for genes known to be involved with wing patterning. These genes included several known *B. anynana* wing patterning genes, such as those coding the proteins Antennapedia (*antp*; Matsuoka & Monteiro, 2019; Saenko, Marialva, & Beldade, 2011), apterous (*ap*; Prakash & Monteiro, 2018), CD63 antigen (*CD63*; Özsu, Chan, Chen, Gupta, & Monteiro, 2017), Cubitus interruptus (*Ci*; Monteiro & Prudic, 2010), decapentaplegic (*dpp*; Connahs et al., 2019), Distal-less (*Dll*; Monteiro et al., 2013, doublesex (*dsx*; Prakash & Monteiro, 2020), Ecdysone Receptor (*EcR*; Bhardwaj et al., 2018), Engrailed (*en*; Monteiro, Glaser, Stockslager, Glansdorp, & Ramos, 2006), hedgehog (*Hh*; Saenko et al., 2011), Invected (*inv*; Monteiro et al., 2006), Notch (*N*; Beldade & Peralta, 2017), patched (*ptc*; Beldade & Peralta, 2017), Spalt (*sal*; Monteiro et al., 2006), Ultrabithorax (*Ubx*; Matsuoka & Monteiro, 2019; Monteiro & Prudic, 2010), and wingless (*wg*; Özsu et al., 2017), as well as genes known to be critical for wing patterning in *Heliconius* and other butterflies, including *aristaless* (Westerman et al., 2018), *BarH-1* (Woronik et al., 2019), *cortex* (Nadeau et al., 2016), *optix* (Reed et al., 2011), and *Wnt* (Martin & Reed, 2010, 2014).

## Results

Sequencing generated over 387 million SE reads (Table S2). Adapter trimming removed 87,301 reads (0.02% of the raw sequenced reads) prior to downstream analysis. Approximately 340 million (87.7%) of the remaining trimmed reads mapped to the *B. anynana* genome. Of the 22,642 annotated genes in the genome, 15,735 (69.5%) were overlapped by at least 10 reads across all libraries and used as the expression set for differential expression analysis. Blast2GO analysis resulted in the functional annotation of 13,498 (59.6%) genes in the *B. anynana* genome, with a total of 40,857 GO terms assigned to genes in the assembly.

A total of 8,448 (53.7% of all expressed genes) genes were differentially expressed between the larva and adult stage heads, with 4,402 upregulated in adult heads and 4,046 upregulated in larva heads (FDR<0.05; Fig. 2; Table S3). GO enrichment analyses found that 248 GO terms were enriched in the heads of adults (Table S4). When reduced to the most specific terms (i.e., parent functions with a significant child GO term were removed to reduce redundancy), 57 enriched GO terms remained, with the top three terms being proton transmembrane transporter activity (FDR=6.21E-10), odorant binding (FDR=1.95E-09), and iron ion binding (FDR=2.41E-09) (Table 1; see Table S5 and Figs. S1-S3 for full results). By contrast, 200 GO terms were enriched in the heads of larvae (Table S6). A total of 47 GO terms remained after reduction, with the top three terms being protein folding (FDR=2.14E-11), structural constituent of ribosome (FDR=6.12E-10), and nucleolus (FDR=8.49E-09) (Table 2; see Table S7 and Figs. S4-S6 or full results).

**Fig. 2:**
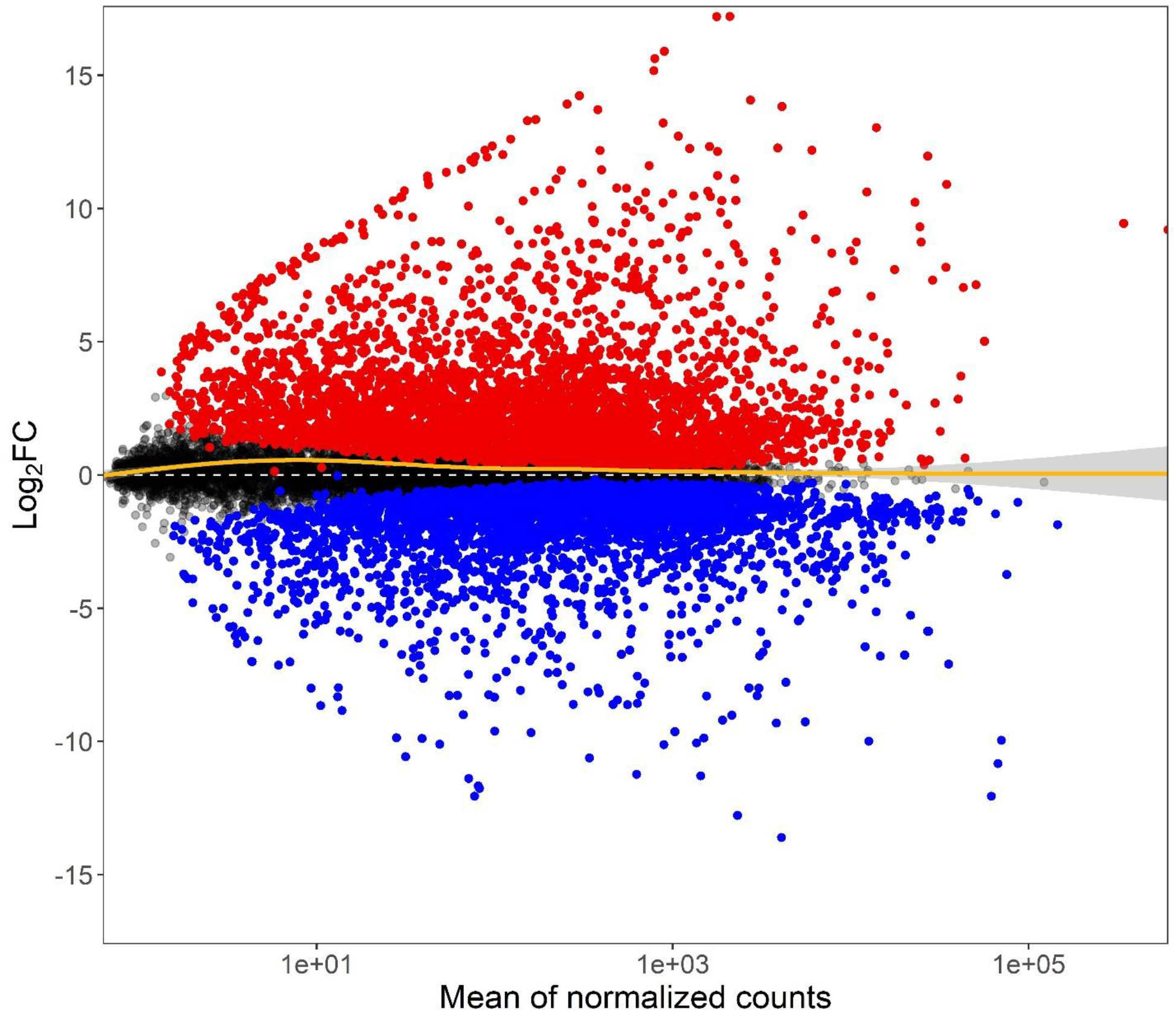
MA plot of the expression level (Mean of normalized counts) and ratio (Log_2_FC) for each gene in *B. anynana* adult heads relative to larval heads. Differentially expressed genes (FDR<0.05) upregulated in adults are represented in red, and those upregulated in larvae are represented in blue. The yellow line is a generalized additive model that was fit to the data.

**Table 1:** Top 10 most specific GO terms enriched in adult heads. BP=Biological Process, MF=Molecular Function, FDR=False Discovery Rate.

| GO ID | GO Name | GO Category | FDR |
| --- | --- | --- | --- |
| GO:0015078 | proton transmembrane transporter activity | MF | 6.21E-10 |
| GO:0005549 | odorant binding | MF | 1.95E-09 |
| GO:0005506 | iron ion binding | MF | 2.41E-09 |
| GO:0020037 | heme binding | MF | 7.36E-09 |
| GO:0071805 | potassium ion transmembrane transport | BP | 1.27E-08 |
| GO:0042302 | structural constituent of cuticle | MF | 1.10E-07 |
| GO:0016705 | oxidoreductase activity, acting on paired donors,<br>with incorporation or reduction of molecular<br>oxygen | MF | 7.22E-07 |
| GO:0004930 | G protein-coupled receptor activity | MF | 9.81E-07 |
| GO:0005249 | voltage-gated potassium channel activity | MF | 6.13E-06 |
| GO:0004984 | olfactory receptor activity | MF | 1.53E-05 |

**Table 2:**
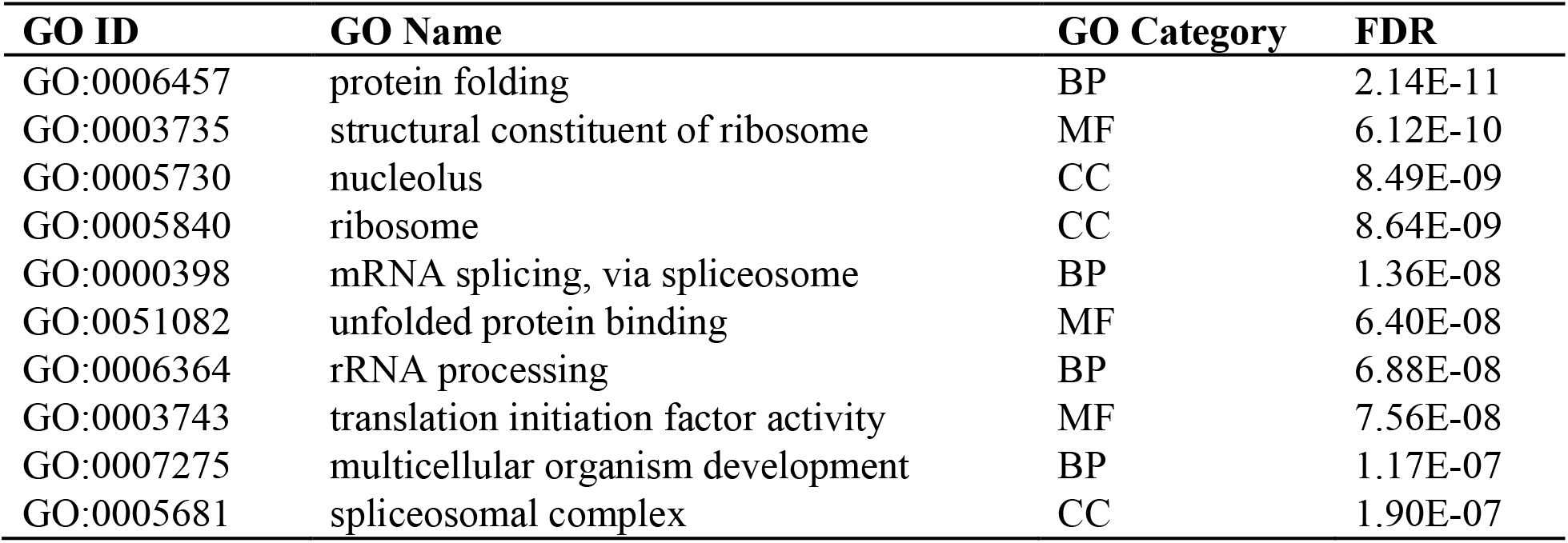
Top 10 most specific GO terms enriched in larval heads. BP=Biological Process, MF=Molecular Function, CC=Cellular Component, FDR=False Discovery Rate.

### Vision genes

Blast hits for 251 of the 274 queried vision genes (92%) resulted in the identification of 1,509 putative homologs in the *B. anynana* expression set (Table S8). Of these expressed homologs, 411 were associated with phototransduction, 70 with eye pigment, and 1,028 with eye development. To identify the top homolog candidates for each of the queried vision genes, we collected the best blast hit, resulting in a set of 251 *B. anynana* vision genes (Table S9). Of these top homologs, 165 (66%) were differentially expressed between larval and adult heads (FDR<0.05), with 83 (53 phototransduction genes, 10 eye pigment genes, and 20 eye development genes) upregulated in adults and 82 (14 phototransduction genes, 5 eye pigment genes, and 63 eye development genes) upregulated in larvae (Fig. 3).

**Fig. 3:**
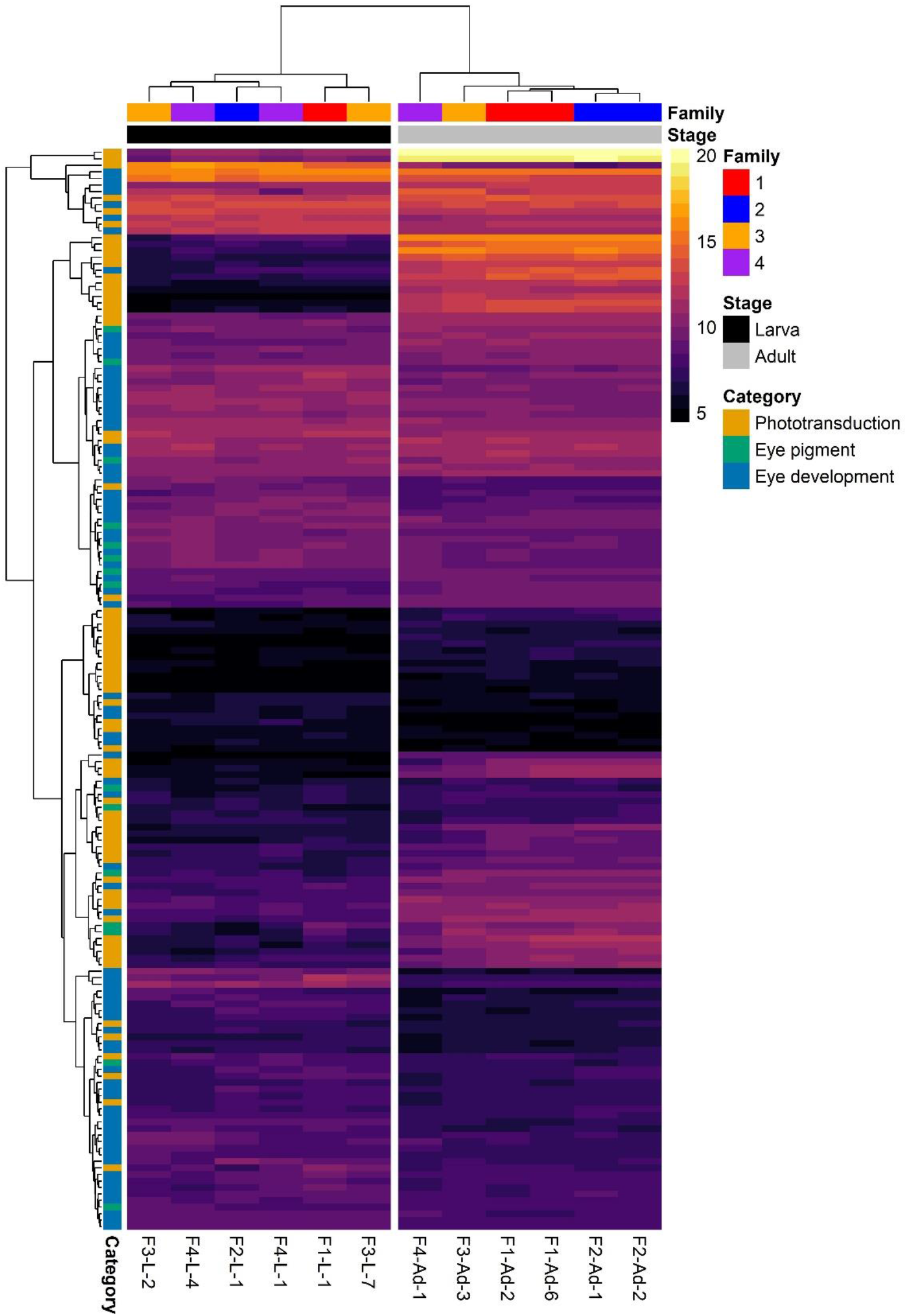
Expression heatmap for differentially expressed top vision homologs in *B. anynana*. Counts were normalized by variance stabilizing transformation, with warmer colors indicating higher expression. Rows denote individual genes, and columns denote samples, both of which are clustered by gene expression. Family indicates the family from which the sample was derived, Stage indicates the developmental stage of the sample, and Category indicates vision gene functions.

A total of 7 opsins (3 visual opsins: *UVRh2, BRh*, and *LWRh*) were identified in the expression set. While all 7 opsins were expressed in both developmental stages, each was significantly upregulated in adults relative to larvae (*UVRh2*, log_2_FC=8.74, FDR=2.93E-219; *BRh*, log_2_FC=7.80, FDR=4.10E-76; *LWRh*, log_2_FC=9.44, FDR< 2.22E-308; *Rh7*, log_2_FC=1.25, FDR=1.35E-4;*pteropsin*, log_2_FC=2.96, FDR=5.77E-4; *unclassified*, log_2_FC=14.1, FDR=1.10E-58; *RGR-like*, log_2_FC=4.94, FDR=3.29E-12; Fig. 4).

**Fig. 4:**
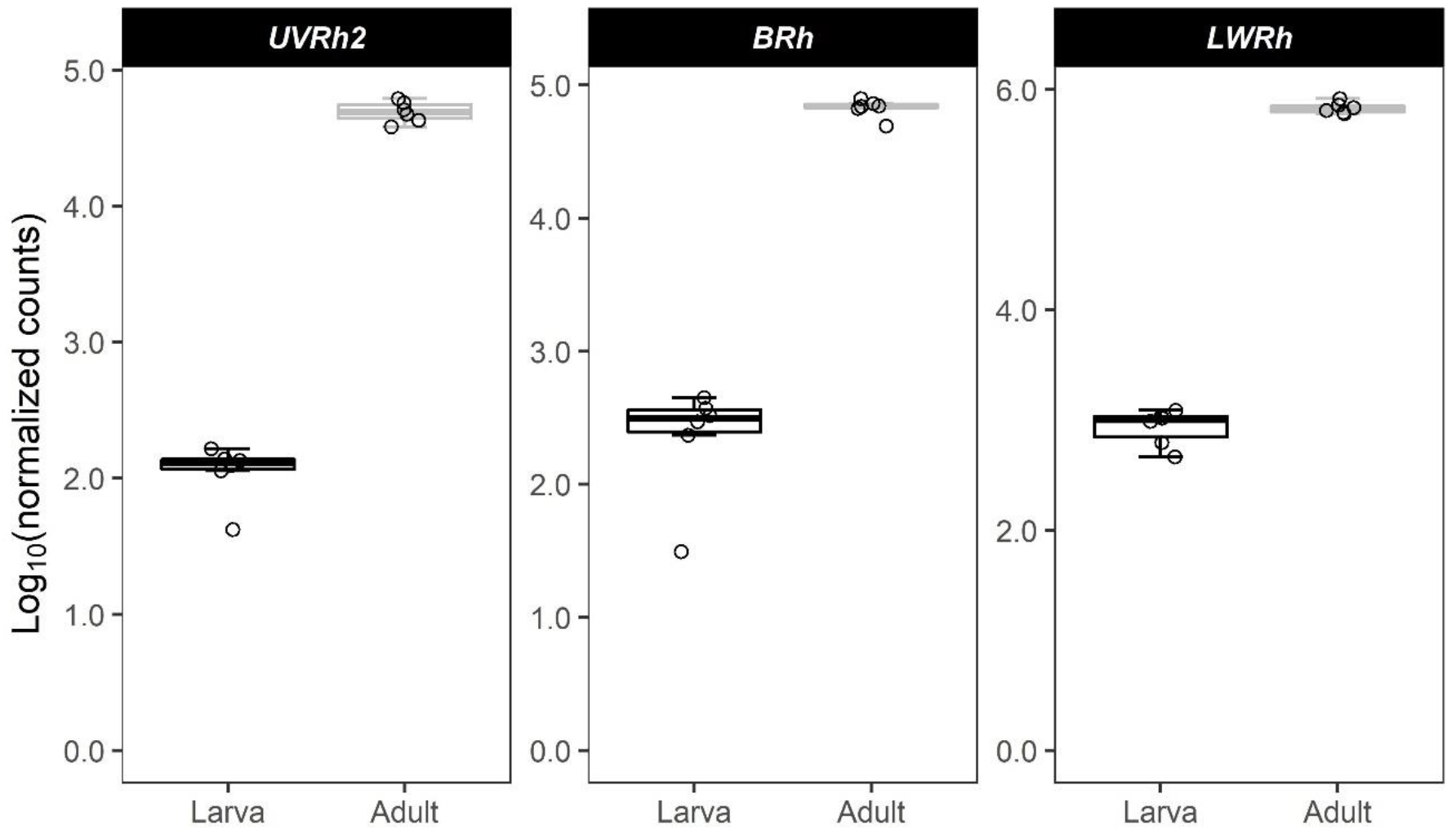
Log transformed normalized counts of visual opsin genes in larvae and adults. Gene labels are *H. melpomene* gene names from Macias-Muñoz et al. (2019). All opsins were expressed in both stages. Horizontal lines within the boxes denote the median. The upper and lower bounds of the boxes indicate the 25th and 75th percentiles, and whiskers extend to the largest count value ≤1.5 × the interquartile range. Y-axes are best fit for each gene.

In addition to the *H. melpomene* and *D. melanogaster* vision homologs we identified, manual searches of the Blast2GO functional annotation identified an additional 21 vision-related genes, including numerous genes associated with phototransduction and eye development (Table S10). Of these genes, 13 were differentially expressed between larval and adult heads, with seven upregulated in adults and six upregulated in larvae (Fig. S7).

### Chemosensory genes

#### Odorant binding proteins

Blast hits for 28 of the 276 queried OBP genes resulted in the identification of 48 putative homologs in the *B. anynana* genome (Table S11). We retained only those containing pfam01395 or smart00708 domains, which resulted in a set of 19 *B. anynana* OBP genes, 17 being expressed in the head (Table S12). Of these homologs, 14 (82%) were differentially expressed (FDR<0.05), 12 of which were upregulated in adults and two of which were upregulated in larvae (Fig. 5). One OBP, a homolog of *Hmel-OBP12* (*BANY.1.2.g14367*), was only expressed in adults (Table 3).

**Fig. 5:**
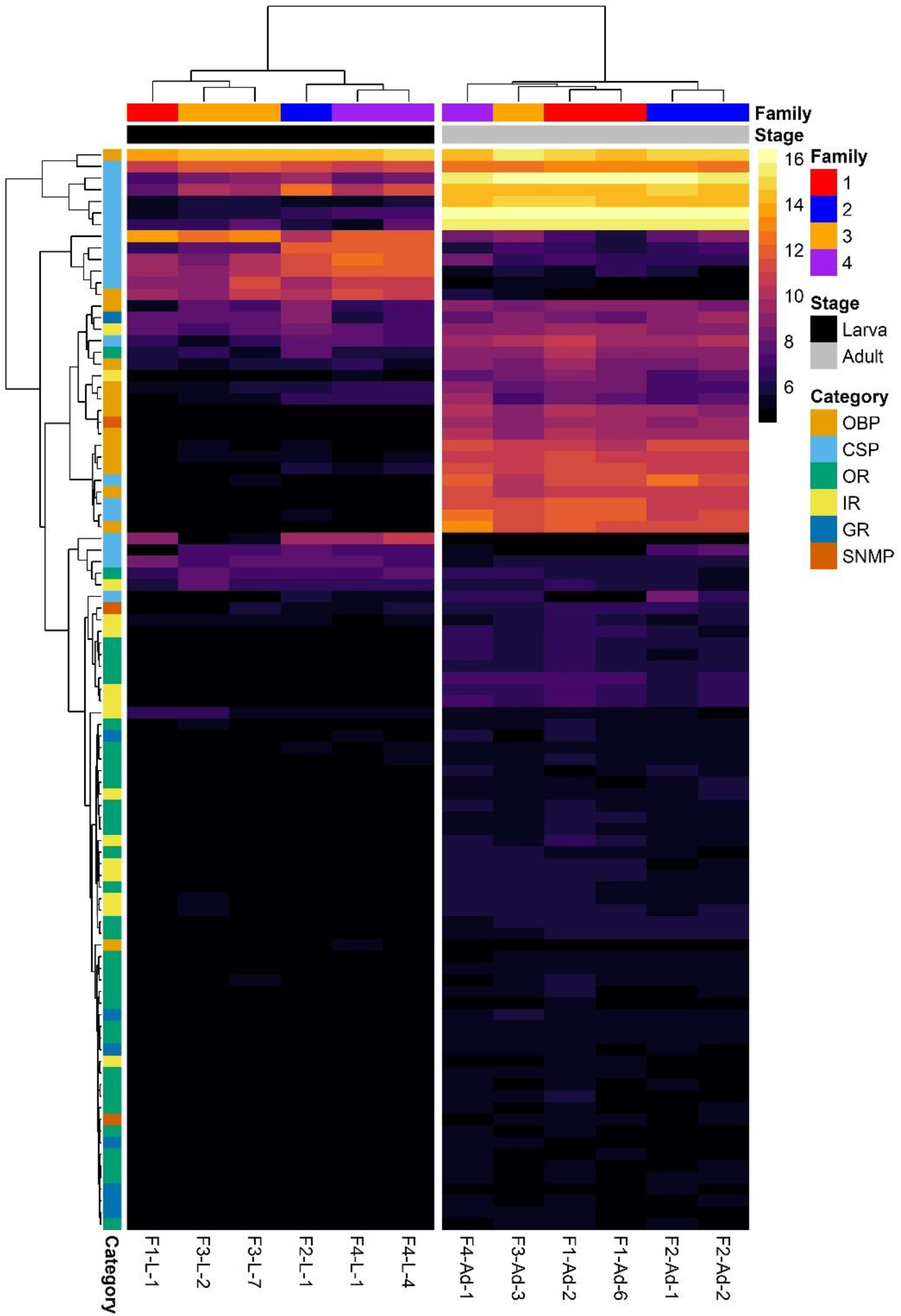
Expression heatmap for differentially expressed top chemosensory homologs in *B. anynana*. Counts were normalized by variance stabilizing transformation, with warmer colors indicating higher expression. Rows denote individual genes, and columns denote samples, both of which are clustered by gene expression. Family indicates the family from which the sample was derived, Stage indicates the developmental stage of the sample, and Category indicates the class of chemosensory gene (OBP=odorant binding protein, CSP=chemosensory protein, OR=olfactory receptor, IR=ionotropic receptor, GR=gustatory receptor, SNMP=sensory neuron membrane protein).

**Table 3:** Chemosensory genes with adult-specific expression. No chemosensory genes were exclusively expressed in larval heads. OBP=odorant binding protein; OR=odorant receptor; IR=ionotropic receptor; GR=gustatory receptor.

| Family | *B. anynana* Gene ID | Query Gene ID |
| --- | --- | --- |
| **OBP** | *BANY.1.2.g14367* | *Hmel-OBP12* |
| **OR** | *BANY.1.2.g00192* | *HmOr3* |
|  | *BANY.1.2.g17040* | *HmOr36* |
|  | *BANY.1.2.g17039* | *HmOr38* |
|  | *BANY.1.2.g02338* | *HmOr41* |
|  | *BANY.1.2.g12362* | *HmOr43* |
|  | *BANY.1.2.g13051* | *HmOr45* |
|  | *BANY.1.2.g23598* | *HmOr45* |
|  | *BANY.1.2.g03909* | *HmOr52* |
|  | *BANY.1.2.g02386* | *HmOr63* |
|  | *BANY.1.2.g22631* | *HmOr69* |
|  | *BANY.1.2.g24109* | *HmOr71* |
|  | *BANY.1.2.g25036* | *HmOr71* |
|  | *BANY.1.2.g24636* | *HmOr74* |
| **IR** | *BANY.1.2.g22328* | *BanyIR31a* |
|  | *BANY.1.2.g20620* | *BanyIR1.2* |
|  | *BANY.1.2.g07365* | *BanyIR40a* |
| **GR** | *BANY.1.2.g00334* | *HmGr5* |
|  | *BANY.1.2.g00335* | *HmGr5* |
|  | *BANY.1.2.g04993* | *HmGr2* |
|  | *BANY.1.2.g15109* | *HmGr19* |

#### Chemosensory proteins

Blast hits for 18 of the 34 queried CSP genes and two genes identified in our manual search resulted in the identification of 27 putative homologs in the *B. anynana* genome (Table S13). Twenty-four of these candidate CSP genes contained the pfam03392 domain, 22 of which were in the head expression set (Table S14). Of these homologs, 19 (86%) were differentially expressed (FDR<0.05), with 11 upregulated in adults and eight upregulated in larvae (Fig. 5).

#### Odorant receptors

Blast hits for 38 of the 70 OR genes and six genes found via a manual search resulted in the identification of 53 putative homologs in the *B. anynana* genome (Table S15). In total, 43 of these were retained as *B. anynana* OR genes by confirmation of the presence of either the pfam02949 or pfam08395 protein domain (Table S16). Of these homologs, 38 were expressed in the head, 34 (89%) of which were differentially expressed between larvae and adults (FDR<0.05). These differentially expressed genes consisted of 33 that were upregulated in adults and one that was upregulated in larvae (Fig. 5). Moreover, 13 ORs were found to exhibit stage-specific expression, all of which were expressed only in adults (Table 3).

#### Ionotropic receptors

We mapped 24 of the 31 *B. anynana* IR sequences from Liu et al. (2018) to genes in the reference genome, 19 of which were expressed in the head (Table S17). Of these homologs, 17 (89%) were differentially expressed (FDR<0.05), 15 of which were upregulated in adults and two of which were upregulated in larvae (Fig. 5). Three IRs were found to only be expressed in adult heads: *BanyIR31a, BanyIR1.2*, and *BanyIR40a* (Table 3).

#### Gustatory receptors

Blast hits for 24 of the 73 GR genes resulted in the identification of 39 putative homologs expressed in *B. anynana* heads (Table S18). We retained only those that contained the pfam08395 domain, resulting in a set of 27 *B. anynana* GR genes, 16 of which were expressed in the head (Table S19). Of these homologs, eight (50%) were differentially expressed (FDR<0.05), all of which were upregulated in adults (Fig. 5). Four GRs, consisting of homologs of *HmGr2, HmGr5*, and *HmGr19*, showed adult-specific expression (Table 3).

#### Sensory neuron membrane proteins

We identified 16 putative SNMP homologs, consisting of blast hits for nine of the 30 SNMP query genes (Table S20). Filtering these putative homologs for genes that contained the pfam01130 domain and were annotated as SNMP genes in the functional annotation resulted in a set of six *B. anynana* SNMP genes (Table S21). All of these were expressed in the head, with three (50%) being differentially expressed between larvae and adults, all of which were upregulated in adults.

### Wing patterning genes

A total of 49 genes associated with wing patterning in butterflies were expressed in the heads of larval and adult *B. anynana* (Table S22). These genes include homologs for *al*, *antp, ap, BarH-1, CD63, Ci, Dll, dpp, dsx, EcR, en, Hh, inv, N, optix, ptc, sal, wg*, and *Wnt*. Of these, a total of 29 (59%) were differentially expressed between larvae and adults (FDR<0.05), including homologs for *al, BarH-1, CD63, Ci, Dll, dpp, en, Hh, inv, N, ptc, wg*, and *Wnt* (Figure S8). Copies of two genes known to be involved with eyespot development in *B. anynana, CD36* and *Ci*, were found to exhibit expression specific to adult and larval heads, respectively.

## Discussion

Our analysis of the gene expression profiles of larval and adult *B. anynana* heads revealed considerable differences between the two developmental stages, with >50% of all expressed genes showing differential expression. Furthermore, we identified numerous genes involved with vision and chemosensation and elucidated how the expression of these genes differs throughout ontogeny. More than 250 *B. anynana* genes putatively linked to vision-related processes were discovered to be expressed in the head, including genes associated with phototransduction as well as eye pigmentation and development. In addition, a total of 118 homologs associated with chemosensation were identified, comprising odorant binding proteins, chemosensory proteins, olfactory receptors, ionotropic receptors, gustatory receptors, and sensory neuron membrane proteins. To our knowledge, this study is the first attempt to characterize the sensory gene repertoire of *Bicyclus* butterflies and provides a promising resource for investigating differences in the sensory biology of larvae and adult butterflies.

### Overall expression differences between larval and adult heads

In larval heads, upregulated genes were linked to developmental processes, including multicellular organism development and the Wnt signaling pathway. Wnt signaling is known to be involved with cell differentiation and proliferation in animals (Logan & Nusse, 2004; Wiese, Nusse, & van Amerongen, 2018). In butterflies, Wnt genes have been found to be involved with wing patterning (Martin & Reed, 2010, 2014) and are expressed in various *B. anynana* tissues during embryogenesis, with *Wnt7* and *Wnt11* both expressed in head tissues (Holzem, Braak, Brattström, McGregor, & Breuker, 2019). In the current study, we found that numerous Wnt genes are expressed in the heads of fifth instar larvae and adults, including *Wnt-1, Wnt-5, Wnt-6, Wnt-10, Wnt-11*, and *WntA* homologs, most of which were upregulated in larval heads.

In addition to developmental processes, genes upregulated in larval heads were also enriched for processes linked to gene expression and protein metabolism. Upregulation of genes involved with these functional categories is possibly in part due to the physiological changes taking place in larval head tissues in preparation for pupation and metamorphosis. As with other holometabolous insects, the larval tissues and organs of butterflies undergo degeneration via autophagy and subsequent remodeling during metamorphosis (see Tettamanti et al., 2008 and Romanelli, Casati, Franzetti, & Tettamanti, 2014 for review). Moreover, in *Manduca sexta*, metamorphic cell death is associated with a marked drop in protein synthesis (Zakeri, Quaglino, Latham, Woo, & Lockshin, 1996). Therefore, the enrichment of GO terms involved with gene expression and protein metabolism might be indicative of a similar decrease in protein synthesis in *B. anynana*.

In adult heads, chemoreception processes were enriched, with associated GO terms including detection of chemical stimulus involved in sensory perception of smell and odorant binding. These results are consistent with elevated chemosensory gene expression in the adult stage of other Lepidoptera (Yang, Yang, Li, Yang, & Zhang, 2016) and suggest a greater investment in the chemosensory system of adult *B. anynana*. Indeed, the adult stage partakes in numerous behaviors that larvae do not, including courtship/copulation, oviposition, and foraging for fruit. Furthermore, *B. anynana* adult females are known to cue in on pheromones produced by males for mate choice (Costanzo & Monteiro, 2007; Nieberding et al., 2008; Westerman & Monteiro, 2013), and chemical cues appear to play just as important of a role as visual cues in mate choice for this species. Finally, the ability of flight permits these butterflies to perform these behaviors in a greatly expanded three-dimensional space compared to the larval stage, possibly requiring adults to maintain a more sensitive and sophisticated chemosensory system.

### Expression of vision-related genes

A large number of vision-related genes were expressed in *B. anynana* heads, most of which were differentially expressed between adults and larvae. These results have implications for furthering our understanding of the differences in the visual capabilities and phototransduction signaling cascade for different life stages of lepidopterans and other holometabolous insects.

The primary visual organs of larval and adult butterflies have disparate morphologies, with larvae possessing two simple eyes consisting of up to six optical units called stemmata and adults having two compound eyes consisting of hundreds of optical units called ommatidia (Ichikawa, 1991). Therefore, it is likely that at least some of the observed patterns in vision-related gene expression in larval and adult heads are due to substantial differences in structure and cell composition. Moreover, during metamorphosis in holometabolous insects, the larval stemmata migrate to the adult optic lobe and continue to function as extraretinal photoreceptors (Briscoe & White, 2005; Gilbert, 1994; Ichikawa, 1991). Consequently, the presence of both adult and larval visual structures in *B. anynana* adults might account for a portion of the upregulation observed in vision genes.

Interestingly, the differentially expressed vision genes upregulated in adult heads were dominated by phototransduction genes (64%), while differentially expressed genes upregulated in larval heads were largely associated with eye development (77%). A greater emphasis on phototransduction in adults is perhaps not surprising, as a significant proportion of the adult head consists of eye tissue, and optic lobes have been found to comprise nearly 75% of the butterfly brain (Ali, 2009; Heinze & Reppert, 2012; Sivinski, 1989). In comparison, the stemmata of larvae occupy a considerably smaller proportion of the larval head. The upregulation of genes involved with eye development in late fifth instar larvae suggests that compound eye developmental processes have initiated just prior to pupation.

Several phototransduction genes were upregulated in larvae relative to adults, including *wunen2*, *ninaC2*, *Cam*, and the innexins *ogre*, *inx2*, and *inx3*. Macias-Muñoz et al. (2019) hypothesize that *wunen* plays a similar role in Lepidoptera phototransduction as *lazaro* in *Drosophila*, which is involved in diacylglycerol (DAG) level regulation (Garcia-Murillas et al., 2006). Lepidoptera have three copies of *wunen*, and while *wunen2* was upregulated in larval *B. anynana*, the other two (*wunen* and *wunen3*) were upregulated in adults. Similarly, *ninaC2* was upregulated in larvae, and an additional copy (*ninaC*) was upregulated in adults. In *Drosophila*, *ninaC* binds to *INAD* and *Cam* to hasten the termination of phototransduction by accelerating arrestin-rhodopsin binding (Liu et al., 2008; Venkatachalam et al., 2010). The observed stage-biased expression of specific *wunen* and *ninaC* copies suggests potential differences in DAG regulation and how the phototransduction cascade is terminated throughout development.

Finally, *ogre, inx2*, and *inx3* form gap junction channels, all of which are critical to visual transmission. Specifically, *ogre* and *inx3* are necessary for visual synaptic transmission in retinal pigment cells in the compound eyes of *Drosophila*, while *inx2* plays an essential role in laminar glial cells (Han, Xiong, Xu, Tian, & Wang, 2017). The fact that these genes are upregulated in larvae suggests the possibility that gap junctions might be either more integral to larval phototransduction or present in greater density in larval eyes. Future functional work should explore this possibility.

Of the visual opsins, *LWRh* had the highest expression in both larvae and adults, which is consistent with previous studies with *B. anynana* (Macias-Munoz et al., 2016) and *H. melpomene* (Macias-Muñoz et al., 2019). This might be a result of *LWRh* expression being in a greater number of photoreceptor cells than *BRh* and *UVRh2*, as *Heliconius LWRh* is expressed in at least six of the nine photoreceptor cells of ommatidia (McCulloch, Osorio, & Briscoe, 2016; McCulloch et al., 2017). It is also interesting to note that all three visual opsins were expressed in larval and adult heads, suggesting that both developmental stages might be capable of perceiving similar wavelengths. Future electrophysiological and behavioral studies should explore the spectral sensitivity and behavioral responses of larval and adult *B. anynana* to different wavelengths of light.

From the current study, it is not possible to determine which of these genes are expressed in the eyes, brain, or other head structures. Numerous studies have localized phototransduction genes in nonvisual tissues, such as the central nervous system (Donohue, Carleton, & Cronin, 2017; Kingston & Cronin, 2015). Therefore, future work should investigate tissue-specific expression and determine the expression patterns of these genes in the eyes. While significant efforts have been made to explore butterfly vision in a number of species, usually focusing on opsins, the results of the current study provide a new set of candidate vision genes for *B. anynana* and will help to expand our understanding of lepidopteran adult, as well as larval, vision.

### Expression of chemosensory-related genes

In the current study, we identified a total of 143 chemosensory genes, most of which showed differential expressed between larval and adult *B. anynana* heads. Notably, we discovered numerous OBPs, CSPs, ORs, IRs, GRs, and SNMPs with either stage-specific expression or displaying differential expression between the developmental stages. Many of these genes share homology with chemosensory genes associated with pheromone detection, host plant recognition, and foraging in other species of Lepidoptera. Because the functions and specificity of chemosensory genes in *B. anynana* are largely unknown, these genes serve as promising targets for further investigation to expand our understanding of chemically-mediated behaviors in this species.

### Odorant binding proteins and chemosensory proteins

Of the various OBPs identified in *B. anynana*, several are of particular interest. The gene *BANY.1.2.g22938* is homologous to *Hmel-OBP13* (alternative name=*HmOBP20*; Dasmahapatra et al., 2012), which was recently found to possibly be involved with species-specific recognition of pheromones in *Heliconius* butterflies (van Schooten et al., 2020). In addition to the two genes sharing sequence homology with *H. melpomene* and *D. plexippus* pheromone binding proteins (PBPs; *BANY.1.2.g06880* and *BANY.1.2.g06881), BANY.1.2.g22938* might also be involved with pheromone detection in *B. anynana*. A homolog of *Eobl-GOBP2, BANY.1.2.g06879*, was also upregulated in adult heads. *Eobl-GOBP2* was previously found to be involved with the detection of plant volatiles in the moth *Ectropis obliqua* (Zhang et al., 2018), suggesting the possibility of a similar role in adult *B. anynana*.

Two OBP (homologs for *Dple-OBP2* and *Dple-OBP19*) and eight CSP (homologs for *HmCSP3*, *HmCSP7*, *HmCSP13*, *HmCSP14*, *HmCSP16*, and *HmCSP17*) genes were upregulated in larval heads relative to adult heads. These genes might be important for larva-specific sensory processing and behavior. Previous studies of OBP and CSP gene expression in larvae and adult *Helicoverpa armigera* noctuidae moths discovered six OBP and four CSP genes that are exclusively expressed in larvae antennae and mouthparts, suggesting that OBP and CSP genes may play a role in larval foraging (Chang et al., 2017). In addition, Zhu et al. (2016) found that larvae of the moth *Plutella xylostella* detect the sex pheromones produced by adults via GOBPs and are attracted to the pheromone in the presence of a food source, possibly allowing larvae to locate promising foraging sites visited by adult females when laying eggs. Thus, it is possible that the OBP genes upregulated in *B. anynana* larvae are involved with foraging or pheromone-specific foraging behavior. Future studies should characterize the function of these genes in larvae to test this hypothesis.

### Odorant, ionotropic, and gustatory receptors

We identified 96 putative chemosensory receptors in the *B. anynana* genome, comprising receptors involved with olfaction and gustation. Thirteen ORs exhibited adult-specific expression, and three pheromone receptor homologs (*HmOR3, HmOR19*, and *HmOR30;* Dasmahapatra et al., 2012; Nakagawa, Sakurai, Nishioka, & Touhara, 2005; Wanner et al., 2007) were upregulated in adult heads, suggesting that they might be involved with mate choice behaviors. Additionally, a homolog of *HmOR49 (BANY.1.2.g06204*), a putative citral receptor (Dasmahapatra et al., 2012; Jordan et al., 2009), was upregulated in adult heads. Citral is released as a male sex pheromone in green-veined white butterfly, *Pieris napi* (Bergström & Lundgren, 1973), and elicits female acceptance behavior during courtship (Andersson, Borg-Karlson, Vongvanich, & Wiklund, 2007). However, as the male pheromones are already characterized for *B. anynana* and do not include citral or its components (Nieberding et al., 2008), it is unlikely that this receptor is involved with pheromone detection. Instead, this receptor might be involved with fruit localization or host plant recognition; citral is also a plant volatile that is present in species such as orange and lemongrass (Martins, Sbaite, Benites, & Maciel, 2011) and serves as a food attractant for *B. mori* larvae (Hamamura & Naito, 1961) and an oviposition deterrent in the light brown apple moth, *Epiphyas postvittana* (Suckling, Karg, Gibb, & Bradley, 1996).

In addition to the OR repertoire, we identified two homologs of *SNMP1* (both *MsexSNMP1* homologs) that were upregulated in adult heads, a protein that forms a complex with pheromone-detecting ORs and an odorant receptor co-receptor (*Orco*; *HmOR2*; identified in the current study as *BANY.1.2.g12855* in *B. anynana*) in insects (Benton et al., 2007; Zhang et al., 2020). *SNMP1* is involved with pheromone detection in both *D. melanogaster* (Benton et al., 2007; Jin, Tal, & Smith, 2008) and numerous lepidopteran species (Krieger et al., 2002; Rogers, Krieger, & Vogt, 2001; Rogers, Steinbrecht, & Vogt, 2001; Rogers, Sun, Lerner, & Vogt, 1997; Zhang et al., 2020). To determine if *SNMP1* might have a similar function in *B. anynana*, studies investigating whether it is expressed in pheromone-responsive neurons, as with other insects (Benton et al., 2007; Forstner et al., 2008; Jiang, Pregitzer, Grosse-Wilde, Breer, & Krieger, 2016), will be necessary. Moreover, future work should investigate whether a similar SNMP1- OR-Orco complex functions in species-specific pheromone detection in *B. anynana*.

Two IRs (*BanyIR75q.1* and *BanyIR93a*) were upregulated in larvae. While the function of *IR75q.1* has not yet been characterized, it belongs to the antennal IR subfamily and is a member of the *IR75* clade expansion (Liu et al., 2018). However, in *D. melanogaster, IR93a* works in combination with *IR21a* and *IR25a* in thermosensation (Enjin et al., 2016; Knecht et al., 2016) and with *IR25a, IR40a*, and *IR68a* for hygrosensation (Knecht et al., 2017, 2016). If *BanyIR93a* functions similarly in *B. anynana*, upregulation of *BanyIR93a* in larvae might suggest the importance of temperature and humidity cues for larval behavior, possibly allowing them to choose an optimal pupation site to avoid desiccation. Interestingly, two IRs upregulated in adults, *BanyIR1.2* and *BanyIR75d*, were found to be putatively involved with seeking host plants for oviposition, as they were upregulated in antennae of females of *H. armigera* that had been mated (Liu et al., 2018). In *B. anynana*, the expression of *IR1.2* was specific to adults, consistent with a possible function in oviposition-related behaviors in this species.

We identified six *B. anynana* GRs (*BANY.1.2.g00334, BANY.1.2.g00335, BANY.1.2.g00336, BANY.1.2.g00338, BANY.1.2.g00341*, and *BANY.1.2.g00343*) homologous to *H. melpomene* sugar receptors *HmGr4*, *HmGr5*, *HmGr6*, *HmGr45*, and *HmGr52* (Briscoe et al., 2013), four of which were expressed in the head. Moreover, three CO2 receptors (*BANY.1.2.g11581, BANY.1.2.g04993*, and *BANY.1.2.g10587*) homologous to *HmGr1, HmGr2*, and *HmGr3* were present in the *B. anynana*, all of which were upregulated in adult heads.

Interestingly, we found four GRs in the *B. anynana* reference genome that are homologous to *HmGr9* and *HmGr57*, two of which were expressed in *B. anynana* heads (*BANY.1.2.g03939* and *BANY.1.2.g11125*). These *H. melpomene* GRs were characterized as putatively being involved with host plant identification via recognition of the plant alkaloid synephrine (Briscoe et al., 2013; van Schooten et al., 2020). While we cannot determine the selectivity of the two *B. anynana* homologs from the current study, it is possible that they might also be involved with host plant recognition in adults and/or larvae. Alternatively, these GRs might be involved with detection of *B. anynana’*s adult food source, ripe/rotting fruit, as synephrine is also present in citrus fruits (Stewart, Newhall, & Edwards, 1964). Regardless, the function of these GRs warrants future investigation.

### Expression of wing patterning genes

Wing patterning genes have been hypothesized to underlie assortative mating behaviors and ultimately speciation in Lepidoptera through associations with preference for the traits they influence (Kronforst et al., 2006; Merrill et al., 2019). This might occur in two ways: (1) both the trait and preference are controlled by the same gene; or (2) the genes controlling the trait and preference for that trait are separate but maintained in high linkage disequilibrium (i.e., inherited together) (Servedio, 2009; Smadja & Butlin, 2009; Westerman, 2019). However, empirical evidence for either of these hypotheses, or for the genetic basis of assortative mate preference more broadly, is relatively slim.

*Bicyclus* is a promising genus to test these hypotheses, as it is very speciose (80+ known species), with many species living in sympatry (Condamin, 1973; Kodandaramaiah et al., 2010). Different species vary in eyespot number and size, potentially permitting the recognition of conspecifics and facilitating assortative mating. Here, we found that numerous genes known to be involved with wing patterning in butterflies were expressed in *B. anynana* heads, possibly in the brain, eyes, or both tissues. If these genes, particularly those involved with eyespot development in *B. anynana*, are linked to preferences for eyespot traits, they might play a role in the great amount of diversity we see in this taxon. We propose that these wing patterning genes should be investigated as potential drivers of assortative mate preference and speciation in *Bicyclus* butterflies.

## Conclusions

In this study, we identified the sensory gene repertoire of the butterfly *B. anynana* and characterized the expression of these genes in larval and adult heads. While visual and chemosensory genes have been explored in many adult Lepidoptera, few studies have investigated the expression of such genes in their larval stages. Our results provide an initial step in elucidating the differences in sensory processing throughout development in butterflies. Moreover, we identified numerous candidate genes for host plant recognition, foraging, and mate choice, including both chemosensory and wing patterning genes expressed in *B. anynana* heads. Future studies should explore the function of these candidate genes and determine their tissue specificity.

## Acknowledgements

We would like to thank Jacob Siebenmorgen for help with starting *B. anynana* families for this experiment, Elizabeth Ruck, Amanda Scholes, and Tara Stuecker for their advice on molecular techniques, and Grace Hirzel, Matt Murphy, Sushant Potdar, and Nikki Robertson for assistance with butterfly husbandry. This research was funded by the University of Arkansas and NSF IOS grant #1937201 to ELW.

## Data Accessibility

All sequence data associated with this study are accessible through the NCBI BioProject database (accession ###), including the raw sequence data (Sequence Read Archive accession numbers ###–###). ***To be submitted upon acceptance

## Author Contributions

DAE and ELW designed this research and wrote the paper. DAE performed the research and analyzed the data.

