## Supporting Information for "Ontogenetic changes in sensory gene expression in *Bicyclus anynana* butterflies"

### Supporting Tables

**Table S1:** *B. anynana* sample characteristics.

| | Family 1 | Family 2 | Family 3 | Family 4 | Total |
|---|---|---|---|---|---|
| **# of Larvae** | 1 | 1 | 2 | 2 | 6 |
| **# of Adults** | 2 | 2 | 1 | 1 | 6 |
| **Total** | 3 | 3 | 3 | 3 | 12 |

**Table S2:** Summary statistics for raw sequence data (± SD). Mean mapped reads indicates the number of raw reads that mapped to the *B. anynana* reference genome (v1.2).

| Stage | n | Total raw reads ($\times 10^6$) | Mean raw reads ($\times 10^6$) | Mean mapped reads ($\times 10^6$) |
|---|---|---|---|---|
| Larva | 6 | 187.1 | 31.2 ± 2.9 | 27.1 ± 2.1 |
| Adult | 6 | 200.9 | 33.5 ± 8.5 | 29.6 ± 7.4 |
| All Stages | 12 | 388.0 | 32.3 ± 6.2 | 28.3 ± 5.3 |

**Table S3:** Blast2GO functional annotation of *Bicyclus anynana* genome assembly (v1.2) and DESeq2 results. Genes that were not in the expression set are denoted with “NA.”

**Table S4:** GO enrichment analyses results for genes upregulated in the adult heads.

**Table S5:** GO enrichment analyses results reduced to the most specific terms for genes upregulated in the adult heads.

**Table S6:** GO enrichment analyses results for genes upregulated in the larva heads.

**Table S7:** GO enrichment analyses results reduced to the most specific terms for genes upregulated in the larva heads.

**Table S8:** All putative vision homologs identified in the expression set.

**Table S9:** Top vision homologs (determined by best blast hit) identified in the expression set.

**Table S10:** Additional putative vision-related genes identified via a manual search of the Blast2GO functional annotation.

**Table S11:** All putative odorant binding protein homologs identified in *B. anynana* genome assembly (v1.2).

**Table S12:** Top odorant binding protein homologs (determined by CD-Search) in *B. anynana* genome assembly (v1.2).

**Table S13:** All putative chemosensory protein homologs identified in *B. anynana* genome assembly (v1.2).

**Table S14:** Top chemosensory protein homologs (determined by CD-Search) in *B. anynana* genome assembly (v1.2).

**Table S15:** All putative odorant receptor homologs identified in *B. anynana* genome assembly (v1.2).

**Table S16:** Top odorant receptor homologs (determined by CD-Search) in *B. anynana* genome assembly (v1.2).

**Table S17:** All ionotropic receptor homologs identified in *B. anynana* genome assembly (v1.2).

**Table S18:** All putative gustatory receptor homologs identified in *B. anynana* genome assembly (v1.2).

**Table S19:** Top gustatory receptor homologs (determined by CD-Search) in *B. anynana* genome assembly (v1.2).

**Table S20:** All putative sensory neuron membrane protein homologs identified in *B. anynana* genome assembly (v1.2).

**Table S21:** Top sensory membrane neuron protein homologs (determined by CD-Search) in *B. anynana* genome assembly (v1.2).

**Table S22:** Wing patterning gene homologs identified in the expression set.

### Supporting Figures

**Fig. S1:** Treemap of Biological Process summarizing GO terms upregulated in adults. Enriched GO terms were reduced to the most specific terms in Blast2GO, and redundancy was further reduced using REVIGO. The size of each individual rectangle is relative to the absolute value of $\log_{10}$(FDR) for each GO term. a) All GO term cluster representatives. b) Superclusters of related GO terms.

a

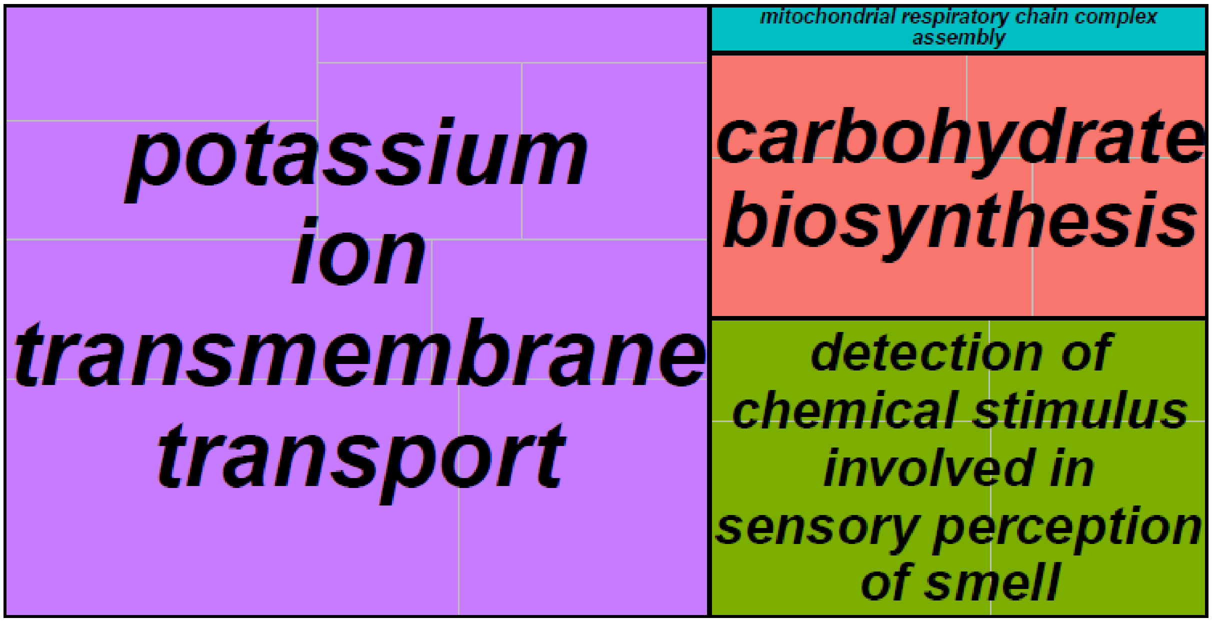


b

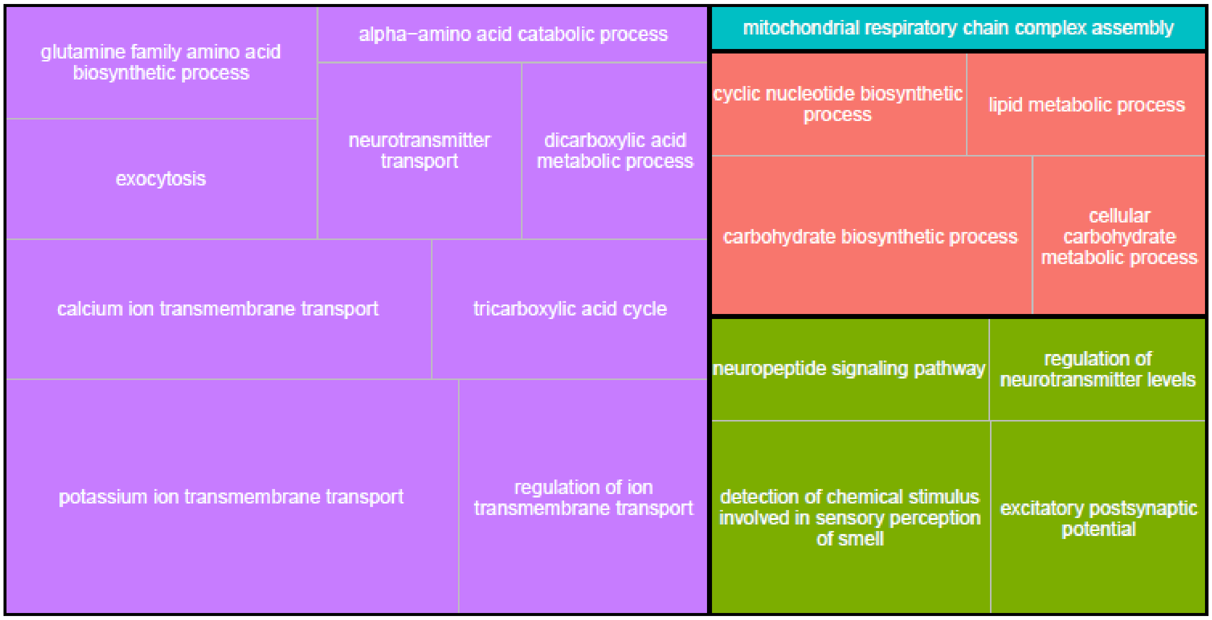

**Fig. S2:** Treemap of Molecular Function summarizing GO terms upregulated in adults. Enriched GO terms were reduced to the most specific terms in Blast2GO, and redundancy was further reduced using REVIGO. The size of each individual rectangle is relative to the absolute value of $\log_{10}$(FDR) for each GO term. a) All GO term cluster representatives. b) Superclusters of related GO terms.

a

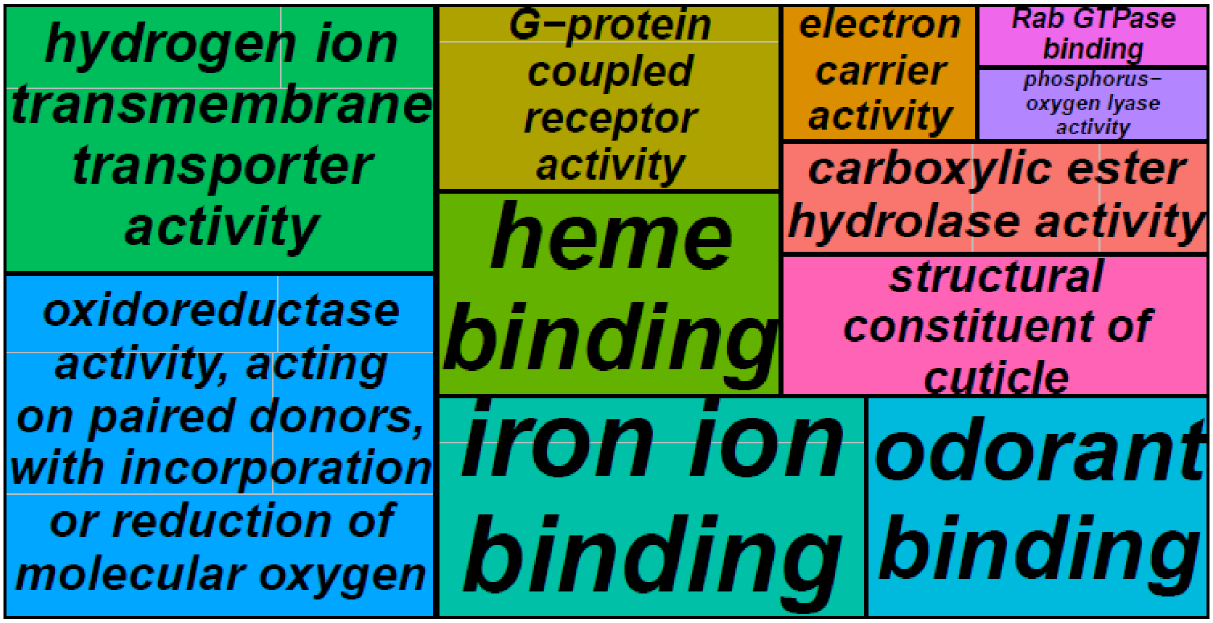


b

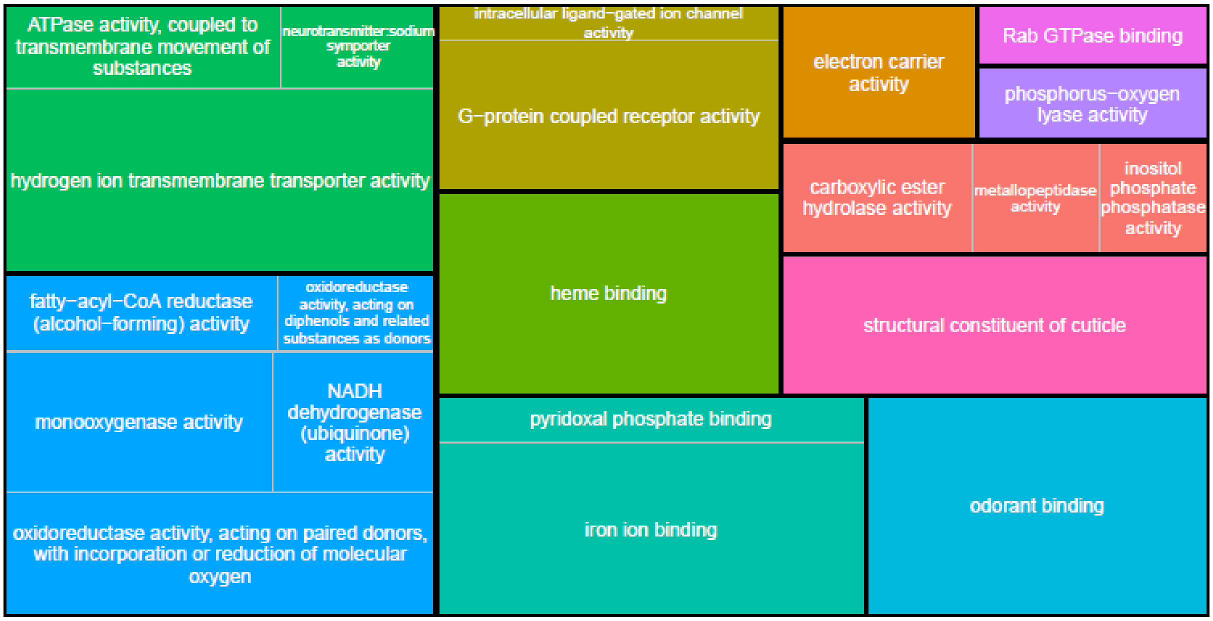

**Fig. S3:** Treemap of Cellular Component summarizing GO terms upregulated in adults. Enriched GO terms were reduced to the most specific terms in Blast2GO, and redundancy was further reduced using REVIGO. The size of each individual rectangle is relative to the absolute value of $\log_{10}$(FDR) for each GO term. a) All GO term cluster representatives. b) Superclusters of related GO terms.

a

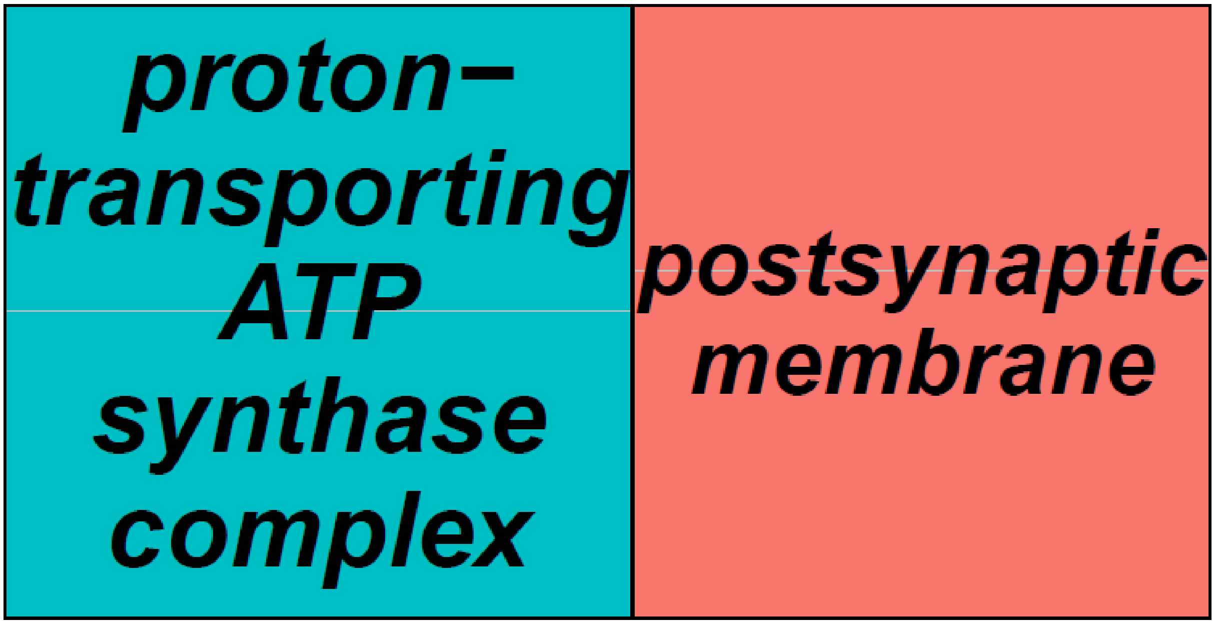


b

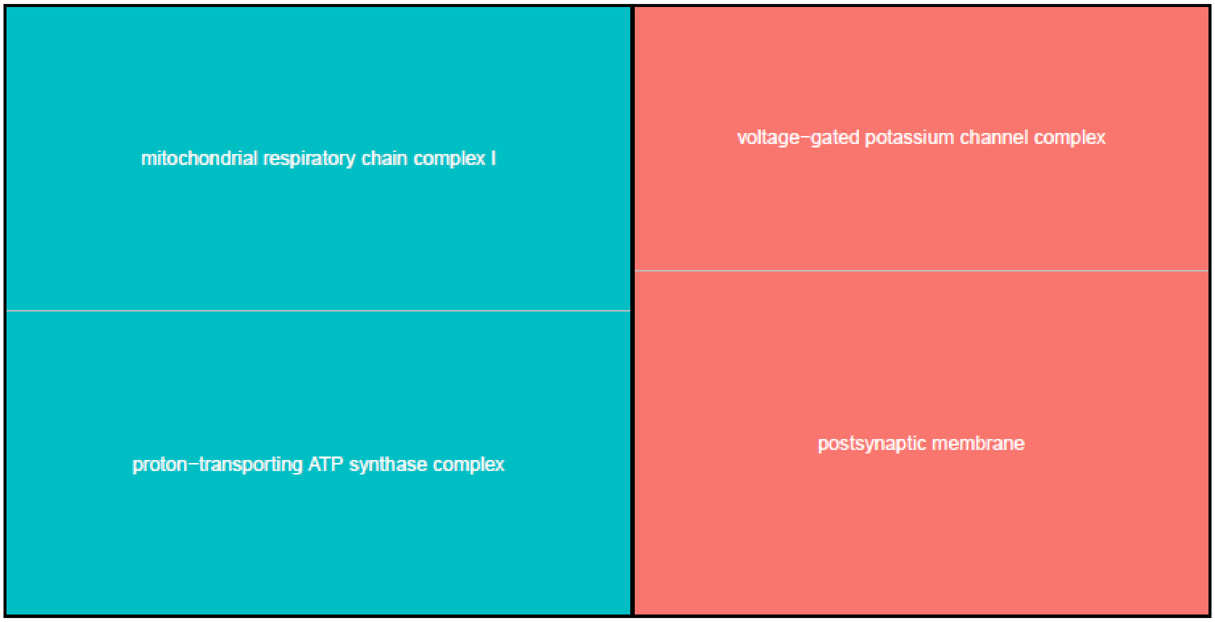

**Fig. S4:** Treemap of Biological Process summarizing GO terms upregulated in larvae. Enriched GO terms were reduced to the most specific terms in Blast2GO, and redundancy was further reduced using REVIGO. The size of each individual rectangle is relative to the absolute value of $\log_{10}(FDR)$ for each GO term. a) All GO term cluster representatives. b) Superclusters of related GO terms.

a

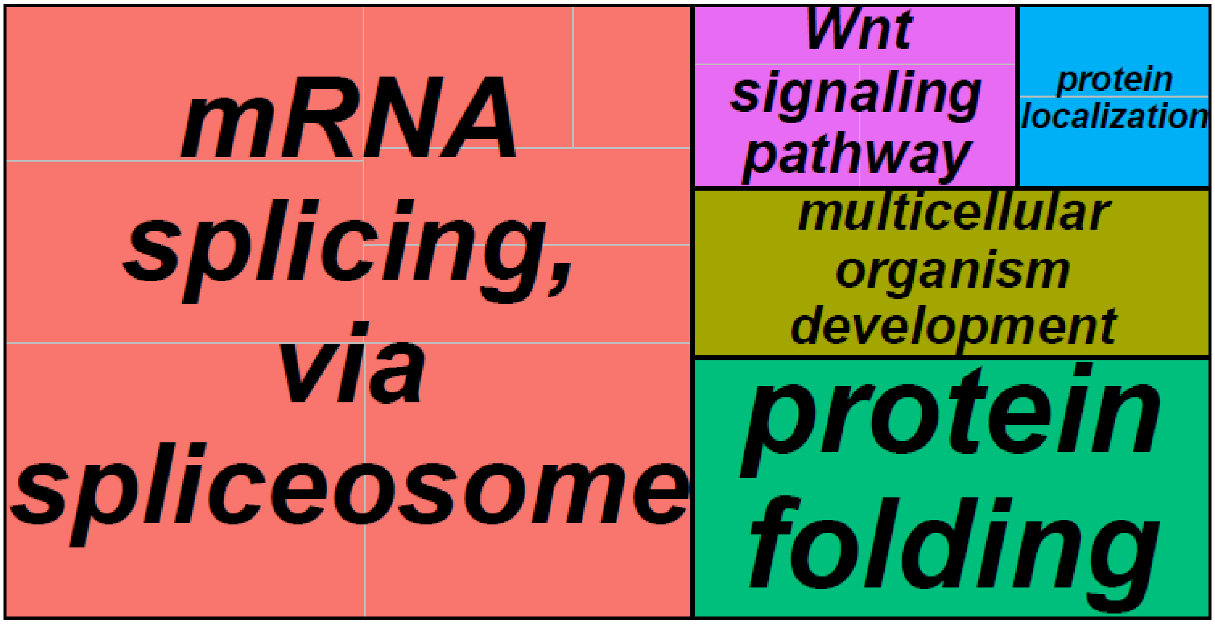


b

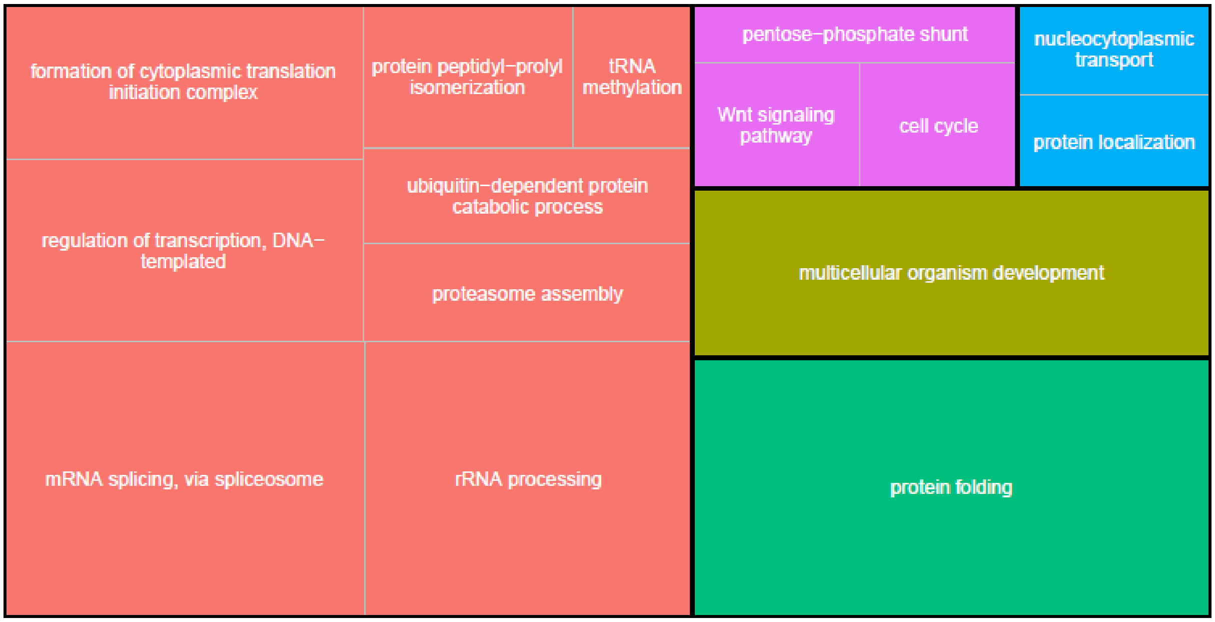

**Fig. S5:** Treemap of Molecular Function summarizing GO terms upregulated in larvae. Enriched GO terms were reduced to the most specific terms in Blast2GO, and redundancy was further reduced using REVIGO. The size of each individual rectangle is relative to the absolute value of $\log_{10}$(FDR) for each GO term. a) All GO term cluster representatives. b) Superclusters of related GO terms.

**a**

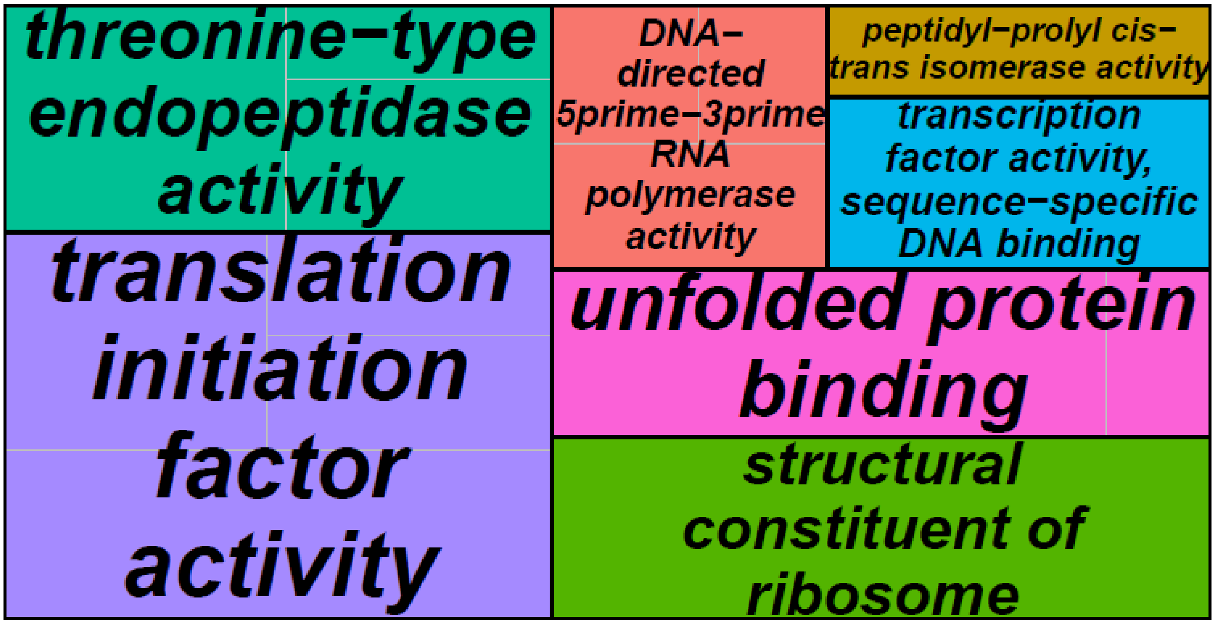


**b**

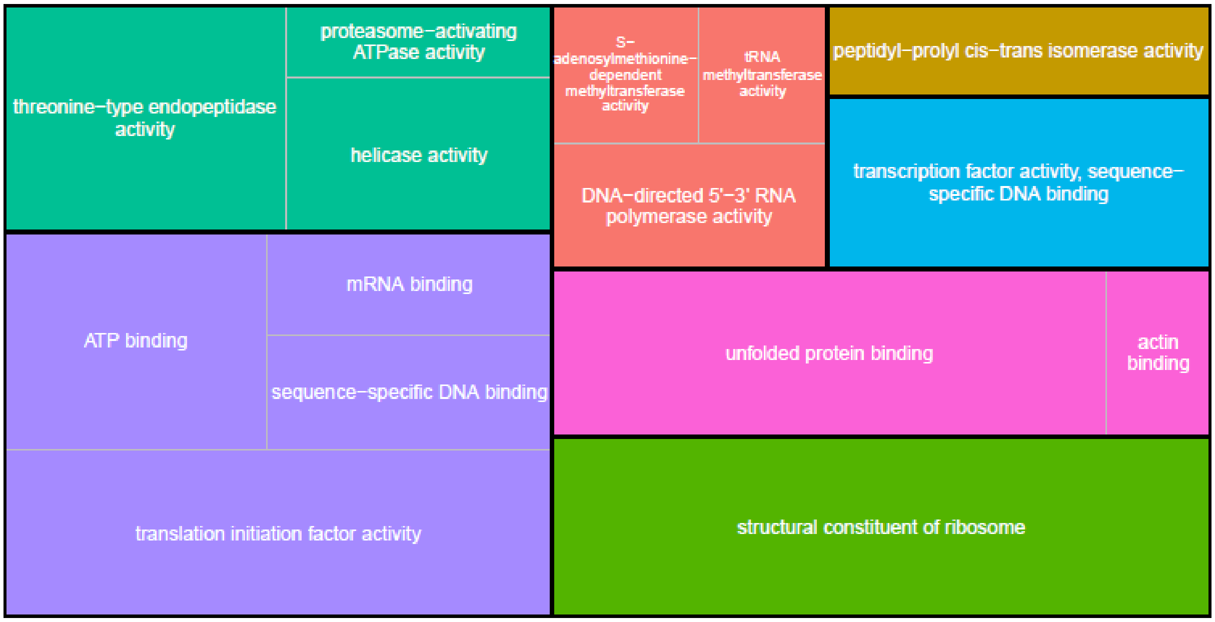

**Fig. S6:** Treemap of Cellular Component summarizing GO terms upregulated in larvae. Enriched GO terms were reduced to the most specific terms in Blast2GO, and redundancy was further reduced using REVIGO. The size of each individual rectangle is relative to the absolute value of $log_{10}$(FDR) for each GO term. a) All GO term cluster representatives. b) Superclusters of related GO terms.

a

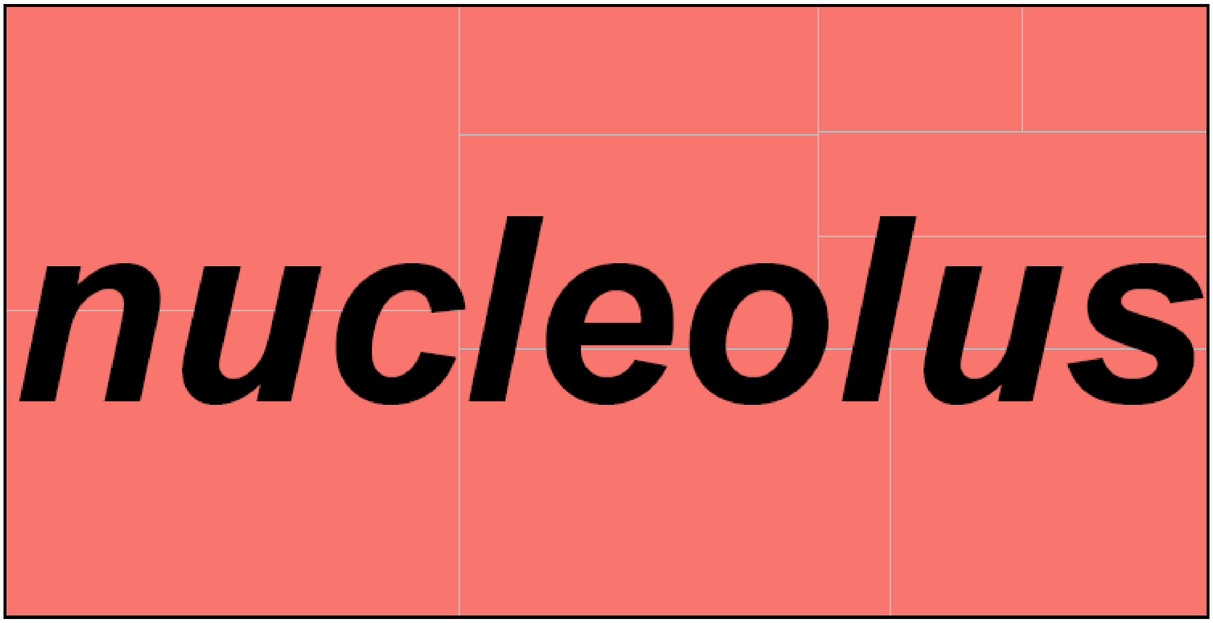


b

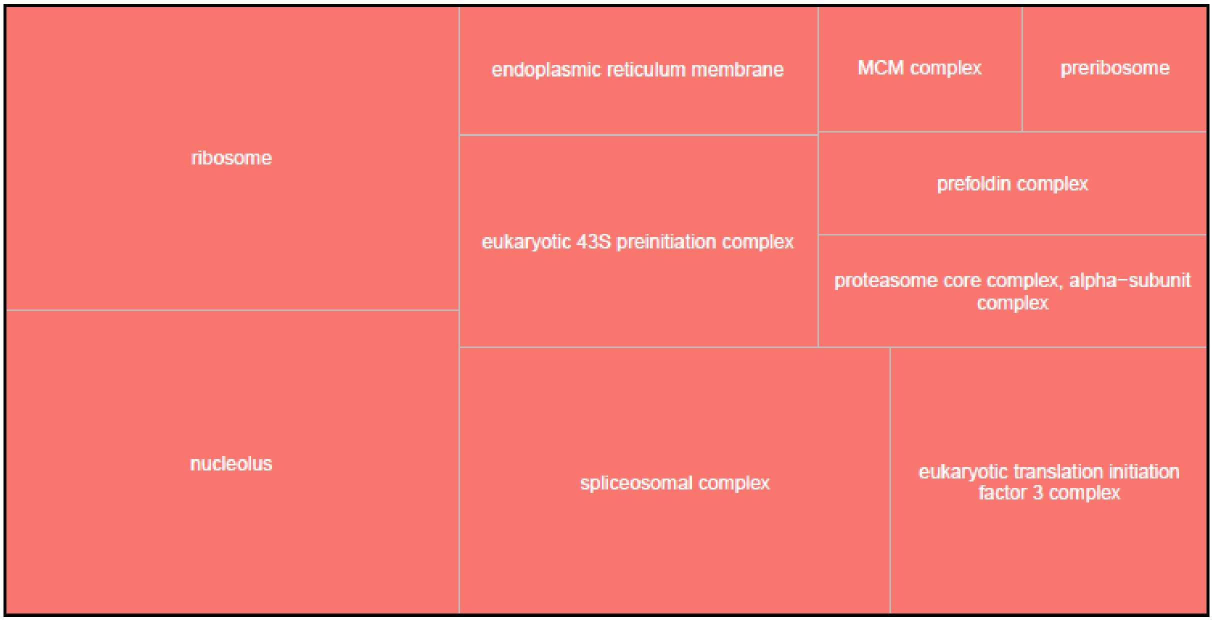

**Fig. S7:** Expression heatmap of additional differentially expressed genes putatively associated with vision. Counts were normalized by variance stabilizing transformation, with warmer colors indicating higher expression. Rows denote individual genes, and columns denote samples, both of which are clustered by gene expression. Family indicates the family from which the sample was derived, and Stage indicates the developmental stage of the sample.

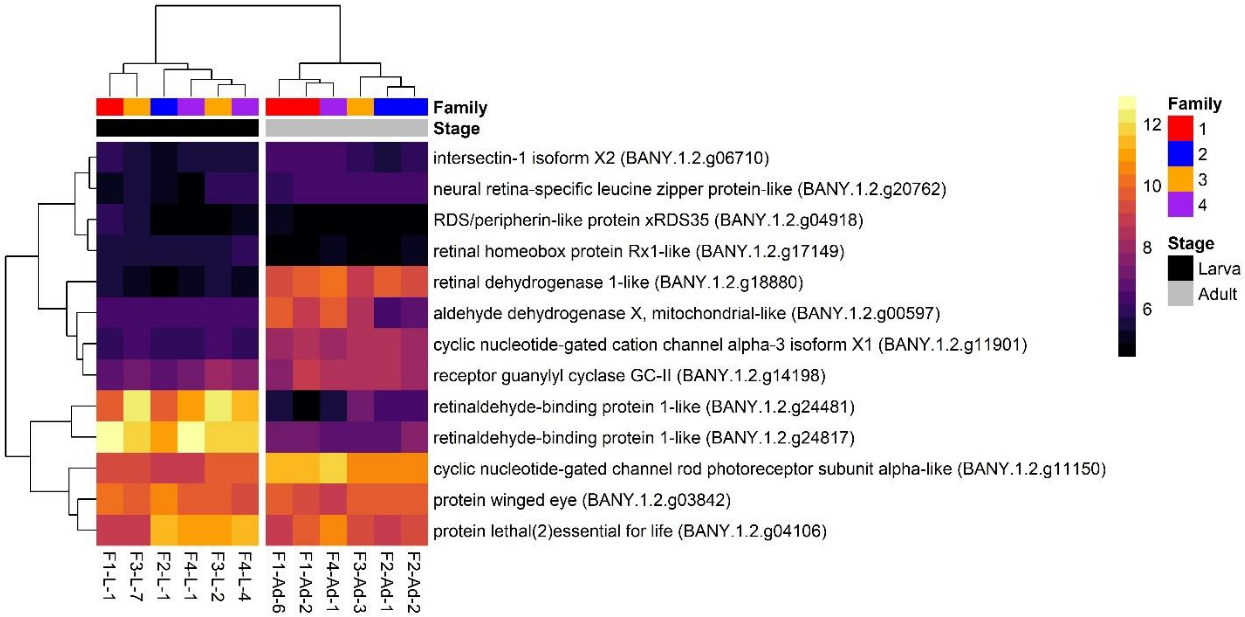

**Fig. S8:** Expression heatmap of differentially expressed genes linked to wing patterning. Counts were normalized by variance stabilizing transformation, with warmer colors indicating higher expression. Rows denote individual genes, and columns denote samples, both of which are clustered by gene expression. Family indicates the family from which the sample was derived, and Stage indicates the developmental stage of the sample. See Table S22 for functional annotation results for each gene.

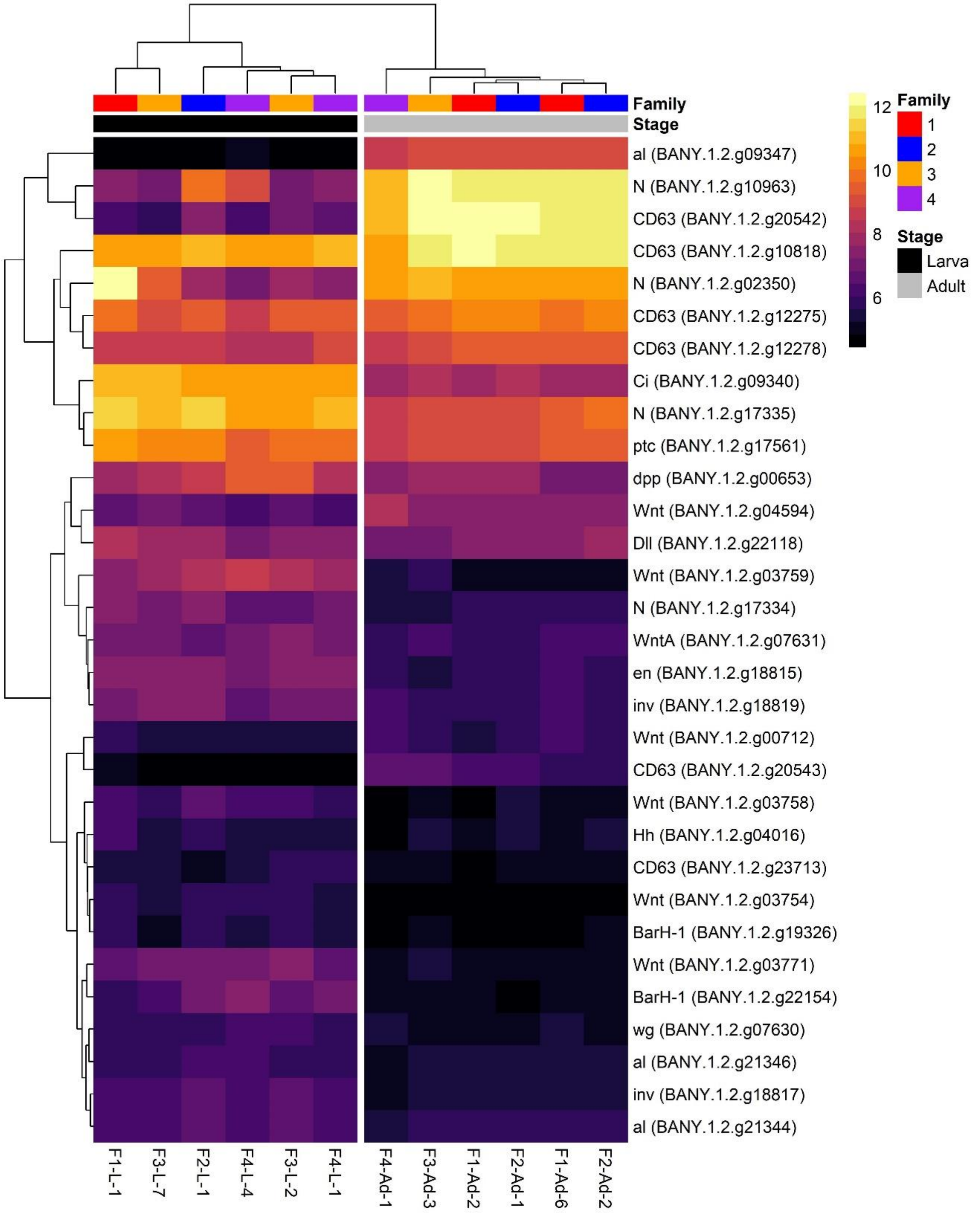
